# TGFβ superfamily signaling regulates the state of human stem cell pluripotency and competency to create telencephalic organoids

**DOI:** 10.1101/2019.12.13.875773

**Authors:** Momoko Watanabe, Jillian R. Haney, Neda Vishlaghi, Felix Turcios, Jessie E. Buth, Wen Gu, Amanda J. Collier, Osvaldo A. Miranda, Di Chen, Shan Sabri, Amander T. Clark, Kathrin Plath, Heather R. Christofk, Michael J. Gandal, Bennett G. Novitch

## Abstract

Telencephalic organoids generated from human pluripotent stem cells (hPSCs) are emerging as an effective system to study the distinct features of the developing human brain and the underlying causes of many neurological disorders. While progress in organoid technology has been steadily advancing, many challenges remain including rampant batch-to-batch and cell line-to-cell line variability and irreproducibility. Here, we demonstrate that a major contributor to successful cortical organoid production is the manner in which hPSCs are maintained prior to differentiation. Optimal results were achieved using fibroblast-feeder-supported hPSCs compared to feeder-independent cells, related to differences in their transcriptomic states. Feeder-supported hPSCs display elevated activation of diverse TGFβ superfamily signaling pathways and increased expression of genes associated with naïve pluripotency. We further identify combinations of TGFβ-related growth factors that are necessary and together sufficient to impart broad telencephalic organoid competency to feeder-free hPSCs and enable reproducible formation of brain structures suitable for disease modeling.

**HIGHLIGHTS:**

- hPSC maintenance conditions influence outcomes in cortical organoid formation
- Identification of an intermediate pluripotency state optimal for cortical organoids
- Feeder support involves activation of diverse TGFβ signaling pathways
- The organoid-promoting effects of feeders can be mimicked by a TGFβ factor mixture

## INTRODUCTION

The unique information processing capacity of the human brain derives from its enormous mass, cellular density, and structural complexity. In general, intellectual ability is proportional to the shape and size of certain brain regions, most notably the neocortex. This structure encompasses neural circuits associated with the execution of motor functions, sensory perception, consciousness, cognition, and language (Roth and Dicke, 2005). Errors in neocortical development can result in a host of defects ranging from gross structural abnormalities to more subtle changes in neural circuit function associated with neurodevelopmental and psychiatric disorders including schizophrenia, autism, and intellectual disabilities such as autism (LaMonica et al., 2012; Nelson and Valakh, 2015). A key to understanding the normal and abnormal functions of the human brain therefore lies in defining the mechanisms regulating neocortical growth, morphogenesis, and circuit assembly.

Traditionally, efforts to model brain development and function have relied on animal models particularly rodents. However, it has become increasingly clear that there are numerous differences between the rodent and human brain (Florio and Huttner, 2014; Hodge et al., 2019), and many human neurological disorders have proven difficult to fully recapitulate (Nestler and Hyman, 2010). The lack of suitable human brain models has also impacted therapeutic discovery, as many drugs discovered to work in animals have failed during clinical trials (van der Worp et al., 2010; Youssef et al., 2018). Human fetal tissue research has been one fruitful approach; however, there are significant ethical and practical concerns including the availability, quality, and heterogeneity of samples. Consequently, a great deal of attention has been placed on the development of alternative methods using human pluripotent stem cells (hPSCs) to create human brain cells and tissues. Over the past few years, several protocols have been established to produce brain organoids, three-dimensional tissue structures that recapitulate the cytoarchitecture organization of the developing brain in vivo (Eiraku et al., 2008; Kadoshima et al., 2013; Lancaster et al., 2013; Pasca et al., 2015; Qian et al., 2016; Watanabe et al., 2017).

While progress in brain organoid technology is rapidly advancing, there are currently no established standards with respect to methodology and quality control. Consequently, marked variabilities in organoid size, organization, and composition can be seen across studies and frequently within experiments. One of the major sources of variability comes from the differentiation protocol, with two major approaches in widespread use: 1) self-patterned whole-brain organoids, in which PSCs are aggregated in a manner that favors spontaneous differentiation and formation of a panoply of both neuronal and non-neuronal cell types (Lancaster et al., 2013; Camp et al., 2015; Quadrato et al., 2017), and 2) region-specific brain organoids, in which the cells are biased or directed towards the formation of single brain compartment such as the cortex or basal ganglia (Kadoshima et al., 2013; Qian et al., 2016; Bagley et al., 2017; Bershteyn et al., 2017; Birey et al., 2017; Watanabe et al., 2017; Velasco et al., 2019; Yoon et al., 2019). Given the limited scope of differentiated cell types formed with the latter approach, there tends to be a higher degree of organoid consistency reflected at the transcriptomic level (Camp et al., 2015; Watanabe et al., 2017; Velasco et al., 2019). However, cytoarchitecture particularly the laminar organization of the cortex can still markedly vary. Moreover, even with a seemingly “reliable” protocol, there can still be differences in the efficiency and overall quality of organoid formation depending on the starting cell lines used as it has been seen in many differentiation protocols (Ortmann and Vallier, 2017).

In our previous work, we established an efficient and reproducible cerebral organoid method that exhibits improved structural integrity including the expansion of the subventricular zone (SVZ), abundant formation of basal progenitors and astrocytes, and the production of upper layer neurons that are localized to the superficial region of the cortical plate (Watanabe et al., 2017). We further documented the similarities in the developmental trajectory of these organoids to the human fetal brain in vivo (Watanabe et al., 2017). Additionally, our methods allow use of developmental patterning signals to create distinct forebrain regions such as the ganglionic eminence (GE), permitting the formation of cortex-GE fusion organoids in which excitatory and inhibitory neurons can intermix to form functional neural networks that mimic activities seen in the human fetal cortex (Samarasinghe et al., 2019). However, in conducting these experiments, we observed that optimal results were only achieved when hPSCs were maintained under particular cell culture conditions, suggesting that even with a well-established organoid protocol, variability may arise from the starting cell population.

PSCs exist in different states: naïve, primed, or a newly proposed transition phase termed formative (Smith, 2017). Naïve and primed states reflect the developmental progression of pre- to post-implantation epiblast populations in vivo with respect to transcriptomic signature, X chromosome dosage compensation, DNA methylation landscape/chromatin state, cell morphology/polarity, cell homogeneity/heterogeneity, metabolism/respiration, and signaling requirements. Formative PSCs are proposed to have intermediate properties of the two extremes, exhibiting features of the early post-implantation embryonic stage in vivo in which lineage-specific gene expression is minimal and cells are still capable of adopting early developmental fates such as a germline stem cell fate (Kalkan et al., 2017; Smith, 2017; Rostovskaya et al., 2019). Naïve and primed PSC states are inter-convertible and metastable in vitro (Weinberger et al., 2016). Perturbation by cell culture conditions can thus alter this balance with subsequent impact on hPSC differentiation capacity and efficiency in vitro.

Here, we define differences in the transcriptional state of hPSCs maintained under different conditions and demonstrate how this in turn relates to success or failure in forebrain organoid development. Organoid formation was optimally achieved when hPSCs were cultured under mouse embryonic fibroblast (MEF) feeder-supported conditions whereas the same hPSCs cultured under feeder-independent conditions yielded poor results. The state of organoid-competency was reversible, as feeder-free hPSCs could efficiently form forebrain organoids when reverted back to MEF-supported conditions. Comparisons of the transcriptome of organoid-competent and -non-competent hPSCs revealed differences in their state of pluripotency, with competency tracking with hPSCs at early stages in the formative transition characterized by expression of multiple genes associated with Transforming Growth Factor Beta (TGFβ) superfamily signaling and naïve pluripotency. Inhibition of TGFβ superfamily signaling in MEF-supported hPSCs impaired efficient formation of forebrain organoids, whereas the addition of 4 TGFβ superfamily growth factors (4G) significantly enhanced forebrain organoid formation from feeder-free cultures. Importantly, 4G supplemented hPSCs exhibited broad competence to form telencephalic organoids resembling cerebral cortex, GE, hippocampus, and choroid plexus. Collectively, these studies identify the state of hPSC pluripotency as a major source of variability in brain organoid formation and illustrate a strategy for modulating this state to help ensure reliability and consistency in organoids to support developmental studies and disease modeling.

## RESULTS

### hPSC growth conditions influence their capacity to effectively form forebrain organoids

To examine how the hPSC maintenance conditions influence forebrain organoid formation, we first compared differentiation outcomes of H9 human embryonic stem cells (hESCs) cultured on particular batches of MEFs and different lot numbers of KnockOut Serum Replacement (KSR) media supplement (hereafter referred to as MEFa and MEFb conditions), or under mTeSR1-based feeder-free conditions on Matrigel (Figures 1A and 1B). When the hESCs reached 70-80 % confluence, cells were dissociated and plated to form one uniformly sized aggregate in each well of a 96-well V-bottom plates (D0) (Figure 1A). After 2.5 weeks in vitro (W2.5), nearly all of the forebrain organoids generated from H9 grown under the MEFa conditions exhibited spherical and epithelialized morphologies with >80% of the cells expressing both FOXG1 and LHX2 along with other forebrain markers (Figures 1C-1G; 1R). Expression of N-CADHERIN (NCAD) and atypical Protein Kinase C (aPKC) was continuous around the periphery of the aggregate, and the apicobasal polarity of neuroepithelial cells was well formed (Figures 1E-1F; 1R). A human induced pluripotent stem cell (hiPSC) line Hips2 and an additional hESC line UCLA1, cultured under the same MEFa conditions, displayed similar efficiency and quality in forebrain organoid formation (Figure S1A-S1E and Watanabe et al., 2017). On the other hand, H9 cultured under the MEFb or feeder-free conditions formed smaller aggregates, with 35% and 15% cells double positive for FOXG1 and LHX2, respectively, and showed poor epithelialized morphology (Figures 1H-1R). Similar results were observed with XFiPSC, a hiPSC line known to yield good organoid results when maintained in the MEFa conditions (Figures 2C-2G and Watanabe et al., 2017).

**Figure 1.**
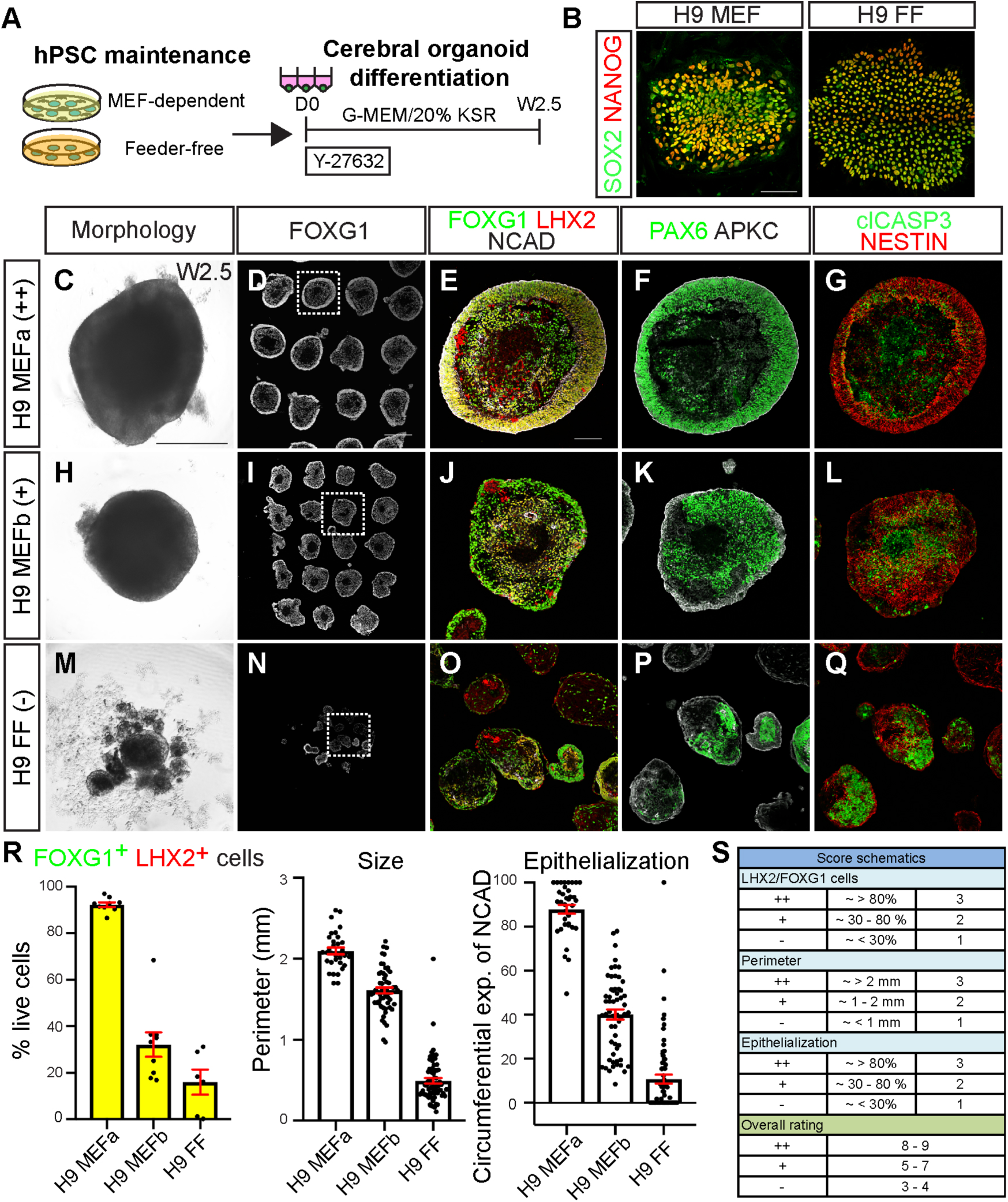
The quality of forebrain organoid formation can vary depending on hPSC maintenance conditions. (A) Schematic of the forebrain organoid protocol from hPSCs maintained under different culture conditions. While the hPSC maintenance conditions were different, the organoid differentiation protocol was identical up to W2.5. (B) H9 hESCs under MEF-dependent or feeder-free conditions display expression of the pluripotency markers SOX2 and NANOG. (C-Q) Assessment of early forebrain organoid differentiation at W2.5. Forebrain organoids derived from H9 maintained with particular MEF batches, MEFa and KSRa (C-G), MEFb and KSRb (H-L), or under mTeSR1-based feeder-free conditions (M-Q). When differentiation was successful, a well-epithelialized layer was observed. Organoids were immunostained for forebrain (FOXG1, LHX2, and PAX6), generic neural (NESTIN), and apical membrane markers (NCAD and APKC). The necrotic core was typically positive for clCASP3. (R) Quantification of percentage of FOXG1^+^ and LHX2^+^ cells out of total live cells per organoid (n = 9 per condition, over 4,400 cells counted per condition), organoid perimeter (n ≥ 32 per condition), and circumferential expression of NCAD defined as the percentage of continuous NCAD expression from a center point (n ≥ 35 per condition). Data are represented as mean ± SEM. (S) Organoid scoring rubric for used to classify forebrain organoid formation at W2.5 as efficient (++), intermediate (+) or poor (-). Scores are based upon the three criteria plotted in (R). hPSCs that formed organoids with a score above 8-9, 5-7, or 3-4, are designated as competent (c-hPSCs), semi-competent (s-PSCs), and non-competent (n-hPSCs). Scale bars: 100 μm (B, E), 500 μm (C, D). See also Figure S1.

**Figure 2.**
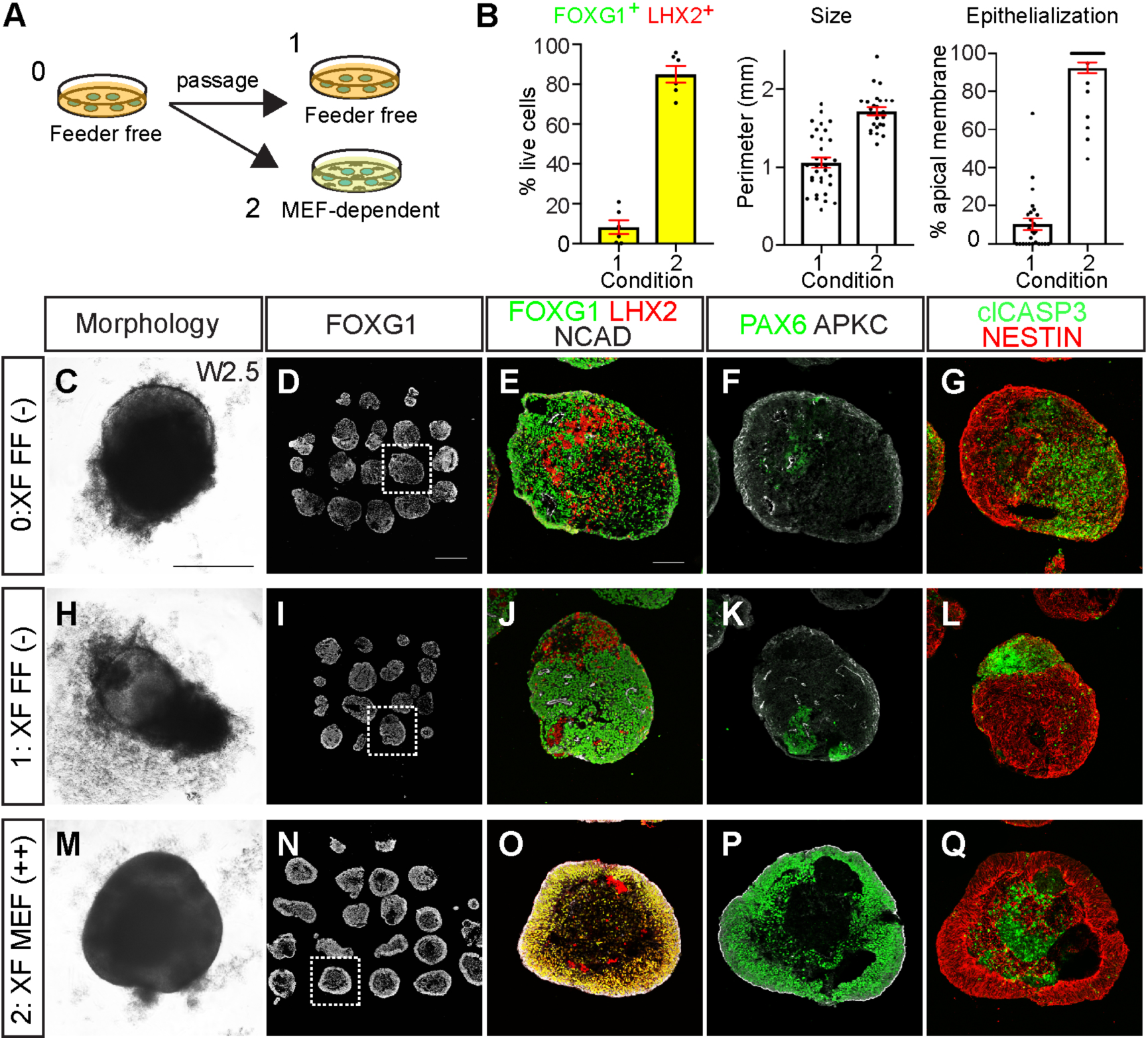
Feeder-free hPSCs regain forebrain organoid competency when passaged under feeder-supported conditions. (A) Schematic depicting how feeder-free hPSCs (condition 0) were either maintained in the same feeder-free culture condition (condition 1) or adapted to MEFa-dependent conditions (condition 2). (B) Quantification of the percentage of FOXG1^+^ and LHX2^+^ cells out of total live cells per organoid (n = 9 organoids per condition; > 6,500 cells counted per condition), organoid perimeter (n ≥ 25 organoids per condition), and circumferential expression of NCAD (n ≥ 26 organoids per condition). Data are represented as mean ± SEM. (C-Q) Morphological and immunochemical analyses of forebrain organoid formation at W2.5. XFiPSCs were initially cultured under feeder-free conditions (C-G, condition 0), and exhibited poor epithelialization and a low percentage of cells positive for cortical (LHX2, PAX6), apical (NCAD, APKC), and generic forebrain (FOXG1, NESTIN) markers. When XFiPSCs were kept under feeder-free conditions (H-L, condition 1), poor organoid formation continued to be observed. However, when XFiPSCs were adapted to MEFa-dependent conditions (M-Q, condition 2), organoid formation was markedly improved. Scale bars: 500 μm (C, D), 100 μm (E). See also Figure S1K-S1T for rehabilitation of another feeder-free line, H9.

To encapsulate the overall success and quality of organoid formation across these and other experiments, we developed a 9-point and three-tier rating scale taking into account three criteria: perimeter size, forebrain marker expression, and extent of epithelialization. High, mixed, and low-quality organoid batches were respectively assigned ratings of ++ (8-9 points), + (5-7 points), and – (≤ 4 points) (Figures 1R and 1S). We hereafter refer to the hPSCs yielding these three outcomes as organoid competent, semi-competent, and non-competent (respectively abbreviated c-hPSCs, s-hPSCs, and n-hPSCs). All the average scores and ratings for different culture conditions and cell lines are summarized in Table S1.

### The states of hPSC forebrain organoid competence are metastable

We next examined if n-hPSCs grown under feeder-free conditions were forevermore fated to yield poor organoid outcomes or could be restored to organoid competence by adapting the cells to the organoid-favorable MEFa culture conditions (Figure 2A). XFiPSCs cultured under feeder-free conditions were accordingly split and passaged in parallel under feeder-free mTeSR1 or MEFa conditions, and then simultaneously differentiated into cortical organoids. In as little as a single passage, XFiPSCs regained organoid competency in stark contrast to the cells that were maintained under the feeder-free conditions (Figures 2B-2Q). The capacity of feeder-free n-hPSCs to be rehabilitated into c-hPSCs by the MEFa conditions was further confirmed using H9 hESCs (Figures S1K-S1T). Together, these results demonstrate that the states of hPSC organoid competency are metastable and can be readily altered by changes in hPSC maintenance conditions.

### c-hPSCs and n-hPSCs exist in distinct transcriptomic states related to TGFβ superfamily signaling and increased expression of genes associated with naïve pluripotency

Given the capacity of hPSCs to switch between states of organoid competence, we sought to define the transcriptomic signatures associated with this difference. We conducted bulk RNA sequencing (RNA-seq) analysis on triplicate samples of both H9 hESCs and XFiPSCs grown under the MEFa and feeder-free conditions. Principal component analyses based on either mRNA transcripts or lncRNAs showed that each hPSC replicate group separated by both cell line origin and cell culture conditions (Figures 3A and S2A). Hierarchical clustering using the top 300 most differentially expressed genes between cell culture conditions revealed that the influences of these conditions were robust (Figure 3B). Gene ontology (GO) analysis demonstrated that genes associated with growth factor signaling, SMAD protein phosphorylation, and developmental processes were highly upregulated in c-hPSCs (Figure 3C). By contrast, genes associated with metabolic pathways such as cholesterol biosynthesis were enriched in n-hPSCs, as has previously been reported for feeder-free cultures (Figure 3C and Garcia-Gonzalo and Izpisua Belmonte, 2008; Zhang et al., 2016; Cornacchia et al., 2019). Extending these GO results, we identified two overt signatures among the genes most highly expressed by c-hPSCs: TGFβ superfamily signaling and genes associated with naïve pluripotency (Figure 3D, and Table S2), whereas some factors associated with primed pluripotency were upregulated in n-hPSCs (Figures 3D, and Table S3). Using Disease Association Protein-Protein Link Evaluator (DAPPLE), a permutation algorithm that looks for significant connectivity in genes associated with a disease state or condition based on known protein-protein interaction data (Rossin et al., 2011), we further confirmed that certain BMP, TGFβ, LEFTY, and NODAL-associated molecules were significantly connected in c-hPSCs (Figure S2C).

**Figure. 3.**
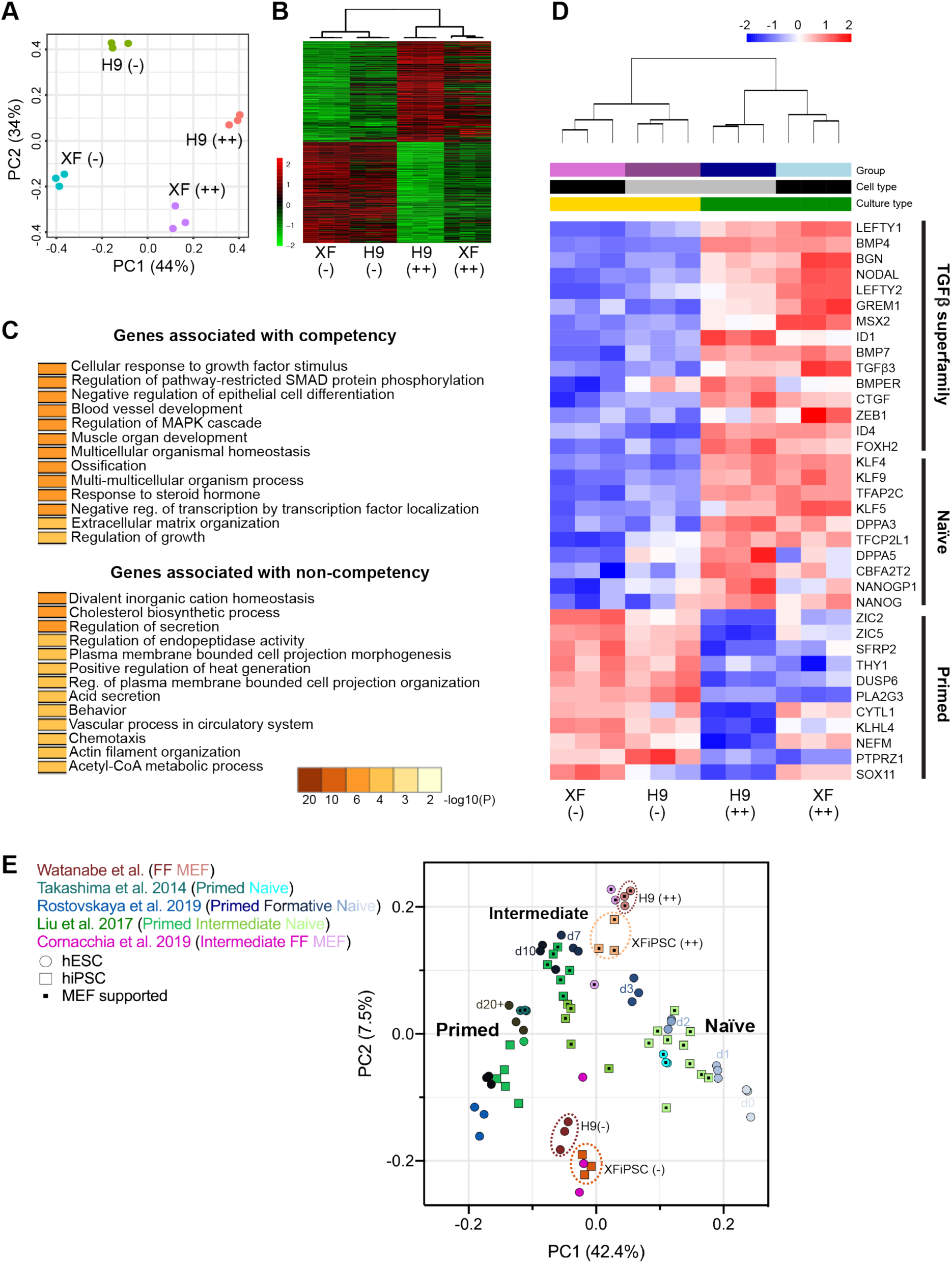
Forebrain organoid competence is associated with a distinct transcriptomic signature defined by elevated TGFβ superfamily signaling and expression of genes connected to naïve pluripotency. (A) Principal component analysis based on RNA transcripts expressed by H9 and XFiPSCs maintained under MEFa-dependent conditions, which is associated with high quality forebrain organoid formation (++), or the same cells grown under feeder-free conditions where poor organoid formation was observed (-). Three independent experimental replicates per cell line and condition are displayed. (B) Hierarchical clustering of the top 300 differentially expressed genes between the samples. The color of the tiles indicates scaled regressed gene expression data. (C) Gene ontology analysis of the upregulated and downregulated transcripts in competent hPSCs, H9 (++) and XF (++), compared to non-competent hPSCs, H9 (-) and XF (-). See also Tables S2 and S3 for the lists of the top 300 genes associated with competency and non-competency. (D) Hierarchical clustering of genes associated with TGFβ superfamily signaling, naïve pluripotency, and primed pluripotency. The color of the tiles indicates scaled regressed gene expression data. (E) Principal component analysis relating the full genomic profile of our H9 and XFiPSC samples to previously published data sets of naïve, formative/intermediate, and primed hPSCs (Takashima et al., 2014; Liu et al., 2017; Cornacchia et al., 2019; Rostovskaya et al., 2019). Days of formative transition from naïve to primed pluripotency (d0-d20+) are indicated according to Rostovskaya et al., 2019. Naïve hPSCs (light green clusters), converted from primed hPSCs or directly from reprogrammed fibroblasts using 5iLAF or 2iLGoY protocol, were previously observed to cluster close to human preimplantation blastocysts in contrast to primed hPSCs grown under E8 media based feeder-free conditions (darker green clusters) (Liu et al., 2017). Some converted hiPSCs, using RSeT or NHSM protocols, from primed hPSCs or directly from reprogramed fibroblasts, showed intermediate transcriptomic signatures (medium green clusters, Liu et al., 2017). Both our MEF-supported and feeder-free hPSCs displayed intermediate signatures, but organoid-competent hPSCs were shifted more toward the naïve hPSC datasets compared to the non-competent hPSCs. See also Figures S2, S3, and Table S4.

As c-hPSCs were prominently associated with MEF-dependent cultured conditions, we further hypothesized that MEF secreted factors must contribute to their transcriptomic state. We accordingly performed mass spectrometry analysis of proteins present in MEF-conditioned culture media (Figure S2D). Among the top 18 proteins identified with greater than 20 spectrum counts, the sixth most abundant protein was Inhibin beta A (INHBA), yet another TGFβ superfamily member whose homodimers constitute ACTIVIN A, a growth factor used in the 5i/L/A method for naïve hPSC conversion (Theunissen et al., 2014). Thus, the strong signature of TGFβ superfamily signaling associated with c-hPSCs likely reflects both autocrine and paracrine actions of the growth factors produced by the hPSCs in addition to signals produced by MEFs such as ACTIVIN A.

We next used cell marker enrichment analyses to compare the transcriptomes of both c-hPSCs and n-hPSCs to reference datasets for naïve and primed hPSCs along with a variety of cell lineages. n-hPSCs were most enriched for genes associated with primed pluripotency with a lower level of association with multi-lineage differentiation (Figures S2B, S2E, and Table S3). Interestingly, while c-hPSCs showed increased expression of naïve pluripotency genes, they did not strongly align with naïve hPSCs in this enrichment analysis (Figures 3D and S2B). At the same time, c-hPSCs also showed a reduced association with multi-lineage differentiation compared to n-hPSCs (Figures S2E and Table S2), suggesting that they constitute a state of pluripotency which is neither primed nor naïve.

To further assess the states of c-hPSC and n-hPSC pluripotency, we compared their transcriptomes to a variety of published hPSC datasets representing both naïve and primed groups, as well as intermediate states representing the formative stages of pluripotency that accompany the developmental transition from pre- to post-implantation blastocysts (Takashima et al., 2014; Liu et al., 2017; Cornacchia et al., 2019; Rostovskaya et al., 2019). Principal component analysis using all genes showed pronounced segregation of naïve and primed hPSC samples, with the latter separating into two groups along the PC1 axis (Figure 3E and S2F). hPSCs representing early to middle stages in formative transition formed clusters in between the naïve and primed states, as previously described (Rostovskaya et al., 2019). When overlaid onto this map of pluripotent stem cell states, our c-hPSCs and n-hPSCs formed distinct groupings, with n-hPSCs positioned among the primed datasets, and c-hPSCs closer to, yet distinct from naïve hPSCs (Figures 3E, S3, and Table S4). The reference datasets that appeared closest to c-hPSCs were those representing days 3 and 7 of formative transition and MEF-supported hESCs recently reported by Cornacchia et al., 2019 (Figures 3E, S3, and Table S4). We further noticed that across all of the datasets, hPSCs that were maintained under MEF-supported conditions were separated from those grown under feeder-free conditions, with the former shifted towards the naïve hPSC state while feeder-free hPSCs tilted towards the primed state (Figures 3E, S3, and Table S4, note the symbols containing or lacking black squares).

Collectively, these data illustrate the wide diversity in the states of pluripotency exhibited by hPSCs maintained under different culture conditions, some of which are favorable and others unfavorable for forebrain organoid formation. Importantly, while our best results were achieved with MEF-supported hPSCs that displayed increased expression of some naïve pluripotency genes, naïve hPSCs themselves yielded poor quality organoids (Figure S1F-S1J). These results are consistent with the recent findings of Rostovskaya et al. who have demonstrated that naïve hPSCs acquire competence for lineage specific-differentiation only after a period of capacitation (Rostovskaya et al., 2019).

### TGFβ superfamily signaling in hPSCs is required for effective forebrain organoid development

Since multiple genes associated with TGFβ superfamily signaling were upregulated in c-hPSCs and ACTIVIN A was being secreted by MEFs, we tested whether TGFβ superfamily signaling was important for forebrain organoid formation. Prior to the start of organoid differentiation, we pre-treated MEF-supported c-hPSCs with either SB-431542, a selective small molecular inhibitor of TGFβ/ ACTIVIN receptors, or LDN-193189, which inhibits BMP receptors at low doses and both BMP and TGFβ/ACTIVIN receptors when used at higher levels (Figure S3A; Inman et al., 2002; Yu et al., 2008). c-hPSCs that were exposed to these inhibitors displayed markedly reduced cortical organoid quality and size, particularly with either low or high doses of LDN-193189 or SB-431542 (Figures S4B-S4C).

Next, we examined whether the addition of TGFβ superfamily molecules to feeder-free n-hPSC cultures could in turn overcome their inability to form high quality organoids. As several TGFβ superfamily molecules such as LEFTYA, BMP4, TGFβ1, TGFβ3, NODAL, and ACTIVIN A, were either enriched in c-hPSCs or produced by the supporting MEFs, we tested the effects of pre-treating n-hPSCs with each of these growth factors alone or in pairwise combinations. Single or double supplementation of most factors, except LEFTYA, showed partial rescue of forebrain organoid formation, with some additive effects (Figure S5, Table S1). The combination of BMP4/TGFβ1 displayed the most potency but was inconsistent in these activities (Figure S5E, Table S1). Moreover, even when there appeared to be favorable outcomes, the overall organoid quality scored below that routinely achieved with MEF-supported c-hPSCs (average score of 5.83 across 6 replicates compared to 8.86 across 7 replicates, Table S1). Taken together, these results demonstrate that BMP and TGFβ superfamily signaling in undifferentiated hPSCs is required for effective cortical organoid production, but individual or pairs of these growth factors are not sufficient to convey organoid competence to n-hPSCs.

### *TFAP2C* and *KLF5* are reliable indicators of hPSC competence for efficient and high-quality forebrain organoid generation

Given that multiple TGFβ superfamily growth factors showed partial benefit to cortical formation, we sought to assess the effects of more complex combinations. However, as our primary readout was quality of organoid formation after W2.5 of differentiation, we needed to employ a more efficient screening strategy. We reasoned that the transcriptional signature of TGFβ superfamily signaling and naïve pluripotency genes that were upregulated in c-hPSCs compared to n-hPSCs could be used to read out the likelihood that the cells would demonstrate success or failure in organoid formation. We thus collected RNA samples from undifferentiated hPSCs cultured under five different conditions: MEFa-dependent XFiPSCs (representing ++ cells), feeder-free H9 supplemented with BMP4 and TGFβ1 (+ cells), XFiPSCs and H9 hESCs grown under mTeSR1-based feeder-free conditions (-), and XFiPSCs grown under E8-based feeder-free conditions (-), and then scored their organoid production quality after W2.5 of differentiation (Figure 4A). Using reverse-transcription quantitative polymerase chain reaction (RT-qPCR) analyses, we found that elevated expression of expression of certain naïve pluripotency-associated genes, *TFAP2C*, *KLF5*, and *DPPA3*, reliably predicted efficient organoid formation more so than genes associated with TGFβ superfamily signaling such as *BMP4*, *TGFβ1 ID4*, *GREM1*, and *ZEB1* (Figure 4B). Moreover, *TFAP2C*, *KLF5*, and *DPPA3* showed very low expression in the hPSCs that yielded the poorest quality organoids (Figure 4B). We further confirmed these results for *TFAP2C* and *KLF5* with additional replicates, cell lines, and cell culture conditions (Figure 4C). While the levels of *TFAP2C* and *KLF5* transcripts were variable, other pluripotency markers such as *OCT4* and *SOX2* were unchanged (Figure 4C). We also observed elevated expression of TFAP2C protein in MEFa-supported H9 hESCs compared to feeder-free H9 hESCs (Figure 4D). Based on these lines of evidence, we conclude that *TFAP2C* and *KLF5* transcript levels can serve as prospective indicators of cortical organoid competence.

**Fig. 4.**
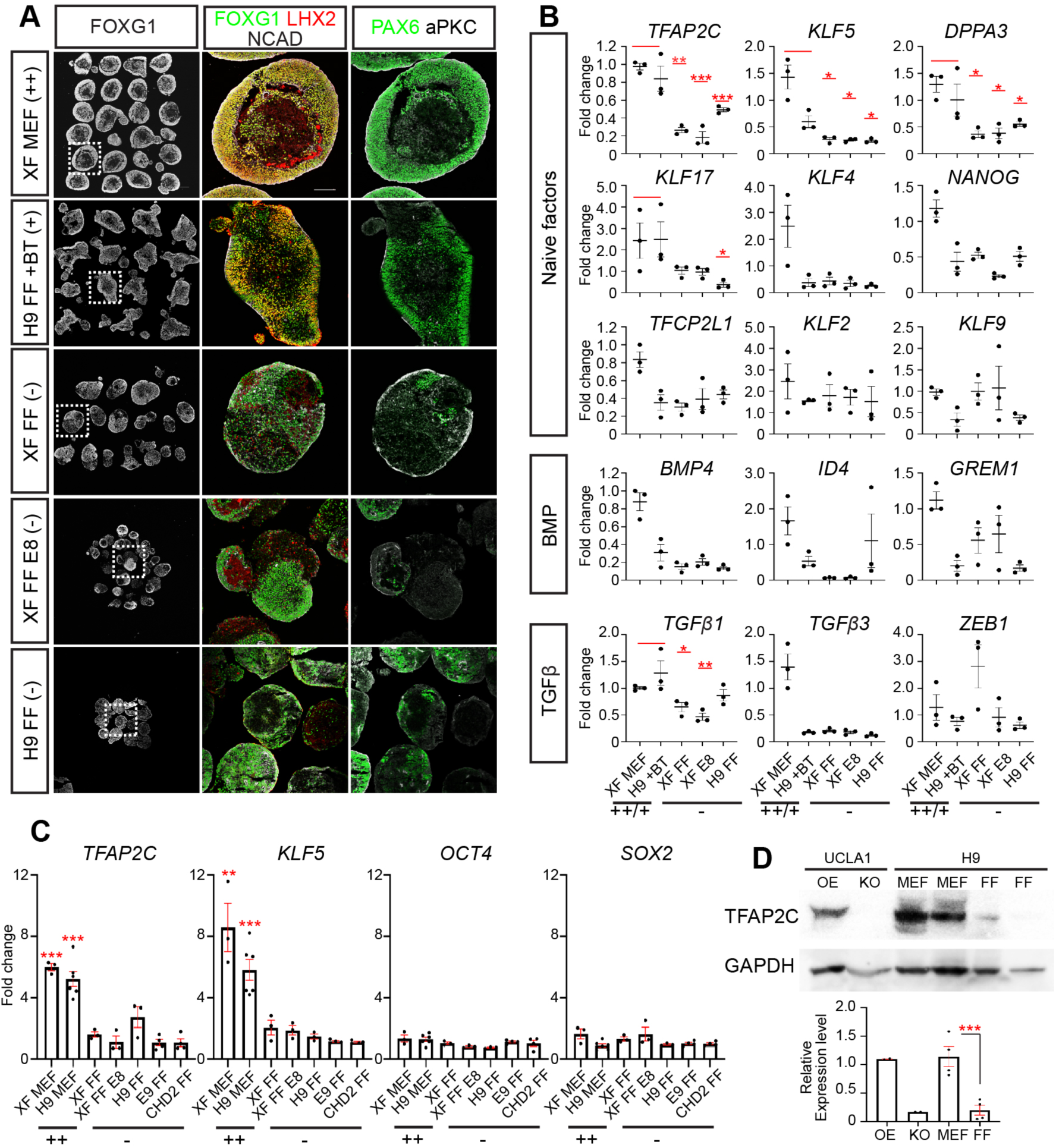
TFAP2C and KLF5 are prospective indicators of hPSCs that can effectively form high-quality forebrain organoids. (A) Representative examples of W2.5 forebrain organoids derived from hPSCs maintained under different culture conditions associated with a range of organoid quality outcomes. Two favorable conditions chosen were XFiPSCs (XF) grown under MEFa-supported conditions (representing ++ cells) and H9 grown under feeder-free conditions supplemented with BMP4 and TGFβ1 (+ cells, see also Figure S5E). Organoid unfavorable conditions included XF grown under mTeSR1-based feeder-free (FF) conditions (- cells), E8-based FF conditions (- cells), and H9 grown under mTeSR1-based FF conditions (- cells). All conditions shown have 3 independent experimental replicates and ≥ 12 organoids per replicate. See also Table S1. (B) RT-qPCR analyses showing that high expression of transcripts associated with naïve pluripotency, *TFAP2C*, *KLF5*, and *DPPA3*, reliably correlated with efficient forebrain organoid development. Expression levels displayed are normalized to XF cultured under the MEFa-supported condition. Data are represented as mean ± SEM. n ≥ 3 independent experimental replicates. Statistical analysis compares the combined two competent conditions XF MEFa (++) and H9 FF + BT (+) against each FF condition, XF FF (-), XF FF E8 (-), or H9 FF (-). (C) *TFAP2C* and *KLF5* expression in additional hPSC lines and replicates, using RT-qPCR analysis. *OCT4* and *SOX2* expression was not significantly different across different cell lines and conditions. The expression levels are normalized to E9 FF hiPSC samples. Data are represented as mean ± SEM. n ≥ 3 independent experimental replicates. Statistical analysis compares E9 FF samples to each condition. (D) Western blot analysis showing high TFAP2C protein expression in H9 grown under MEFa-supported conditions (MEF), compared to H9 FF. Lysates from TFAP2C doxycycline-inducible overexpression line (OE: + dox 1 μg/ml) and knockout lines (KO) were run as positive and negative controls. Relative expression levels normalized to GAPDH are shown. Data are represented as mean ± SEM. n ≥ 4 independent experimental replicates.

### Organoid competence can be conferred onto n-hPSCs by mixture of 4 or more TGFβ superfamily molecules

Using *TFAP2C* and *KLF5* expression as a convenient readout, we systematically tested the impact of pre-treating n-hPSCs with various combinations and concentrations of the 5 TGFβ superfamily growth factors that showed positive benefits in our previous experiments. We included BMP4 and TGFβ1 in all combinations since these two factors had shown the most promising organoid-promoting activities (Figure S5). Eight different combinations of BMP4, TGFβ1, TGFβ3, NODAL, and ACTIVIN A were tested on feeder-free H9 and subjected to RT-qPCR analyses three days after growth factor addition (Figures 5A-5C). Significant increases in *TFAP2C* and *KLF5* without changes in *OCT4* and *SOX2* were observed with three conditions: #7:BMP4/TGFβ1/ACTIVIN A/TGFβ3, #8: BMP4/TGFβ1/ACTIVIN A/NODAL, and #9:all five growth factors together (Figures 5B and 5C). We then carried out forebrain organoid formation from feeder-free H9 cells pre-conditioned with growth factor combinations #4, #7, #8, and #9.

**Figure. 5.**
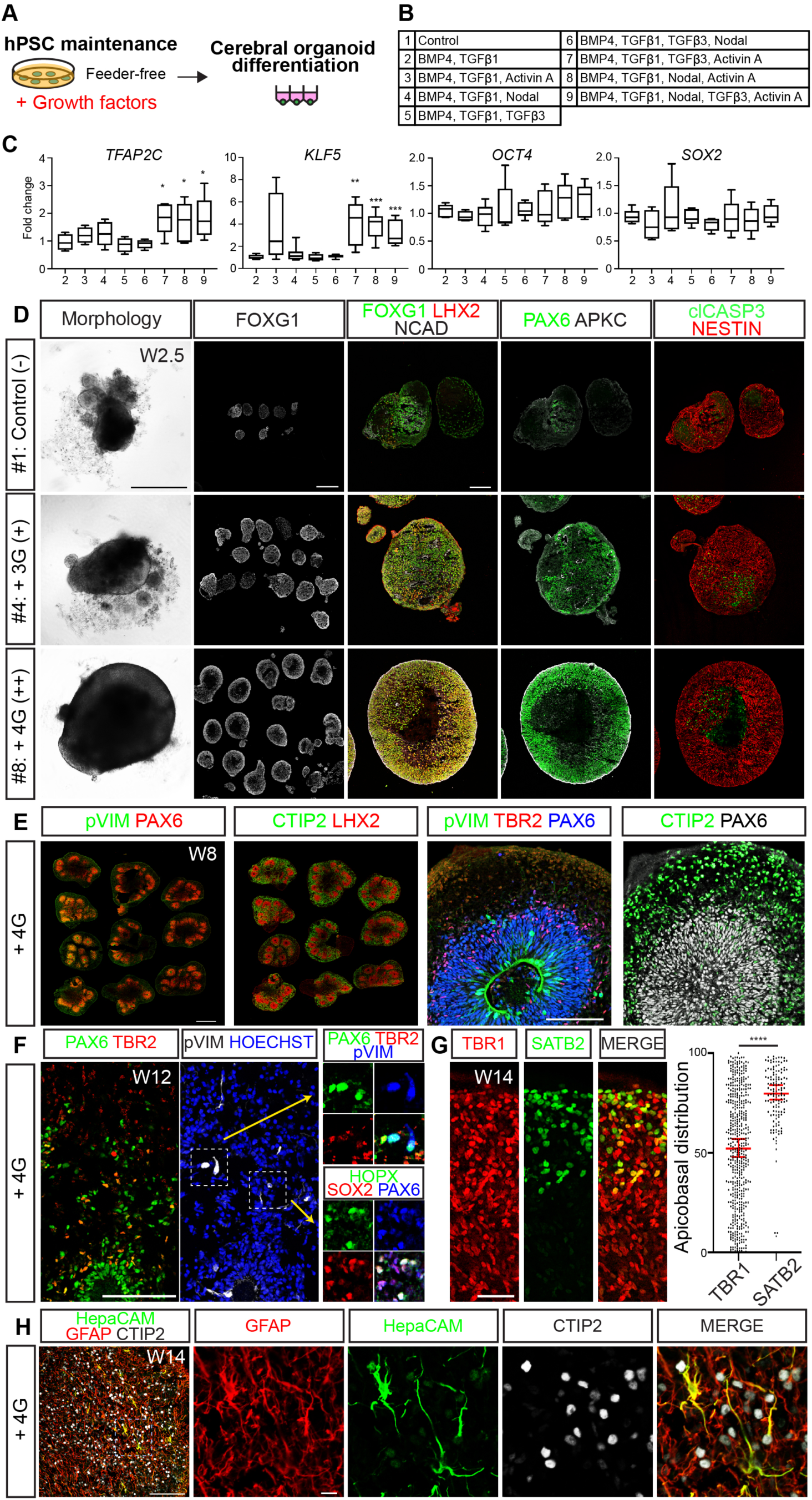
Identification of combinations of TGFβ superfamily growth factors that can enhance forebrain organoid production from feeder-free hPSCs. (A) Schematic of the experimental design. Feeder-free H9 hESCs were supplemented with various combinations of TGFβ superfamily growth factors for three days prior to organoid differentiation. (B) List of the combinations of growth factors tested. (C) RT-qPCR analyses identifying the combinations of TGFβ superfamily growth factors that significantly upregulated the prospective indicators of forebrain organoid competency, *TFAP2C* and *KLF5*. General pluripotency markers, *OCT4* and *SOX2*, were invariant across the different conditions tested. Expression levels displayed are normalized to feeder-free H9 without any growth factor supplementation. Data are represented as mean ± SEM. n ≥ 4 independent experimental replicates. (D) Representative images of forebrain organoids formed from feeder-free H9 supplemented with or without growth factor combinations #4 or #8. Supplementation with four growth factors (4G)-BMP4, TGFβ1, NODAL, and ACTIVIN A (combination #8), enabled efficient and high-quality forebrain organoid induction. (E) Immunohistochemical analysis of 4G-derived cortical organoids at W8. The majority of organoids express cerebral cortex markers (PAX6, LHX2) and exhibit clearly defined laminar architecture reflected in a PAX6^+^ ventricular zone (VZ), TBR2^+^ subventricular zone (SVZ), and CTIP2^+^ cortical plate (CP). pVIM^+^ dividing progenitors were located at the apical membrane of the VZ and in the SVZ region, as seen in the developing cortex. (F) At W12, 4G-derived cortical exhibit an expanded SVZ with abundant formation of basal radial glial cells (SOX2^+^, PAX6^+^, HOPX2^+^, and pVIM^+^). (G) W14 cortical organoids with cells positive for lower-(TBR1) and upper-layer (SATB2) neuronal markers in the CP region. (H) W12 organoids also contain many GFAP^+^ astrocytes along with a smaller number of HepaCAM^+^ cells. Scale bars: 500 μm for morphology, FOXG1, and pVIM PAX6 costaining images in (D); 100 μm for FOXG1 LHX2 NCAD, pVIM TBR2 PAX6, PAX6 TBR2, and HepaCAM GFAP CTIP2 images in (E, F, H); 50 μm for CTIP2 and SATB2 images in (G); and 10 μm for enlarged images in (H).

Without any growth factor supplementation, feeder-free H9 poorly differentiated into forebrain organoids (Figure 5D and Table S1, average quality score of 3.59). Feeder-free H9 that significantly upregulated *TFAP2C* and *KLF5* (supplements #7, 8, 9) all showed a dramatic improvement in organoid formation with quality scores above 7 (Figures 5C-5D, and Table S1). Among these three conditions, the addition of four growth factors (hereafter referred to as “4G”, #7 and #8) yielded the best results (Table S1). Given that our data suggest that the overall level and diversity of TGFβ superfamily signaling matters, we surmise that the addition of 5 factors might result in a signaling profile that is less favorable for forebrain organoid formation. Feeder-free-H9 supplemented with the combination of BMP4/TGFβ1/NODAL (#4), which did not significantly upregulate *TFAP2C* and *KLF5*, showed partial rescue of organoid quality (average score of 5.80, Figure 5D, Table S1), supporting the utility of *TFAP2C* and *KLF5* measurements in predicting organoid outcomes. We confirmed the capacity of the 4G condition #7 to increase *TFAP2C* and *KLF5* expression and enhance cortical organoid formation in feeder-free XFiPSCs and additional two feeder-free n-hiPSC lines (Figure S6), suggesting that the 4G effects may be of general benefit when using feeder-free cells.

In our experience, problems with cortical organoid formation usually becomes manifest within the first few weeks of differentiation, hence our focus on the W2.5 time point throughout this study. However, we wanted to confirm that organoids created from n-hPSCs using the 4G supplementation method exhibited the histological organization and expression of molecular markers of cortical development seen in our previous studies using feeder-supported hPSCs (Watanabe et al., 2017). In our protocol, we add leukemia inhibitory factors (LIF) at W5 to activate STAT3 signaling and thereby stimulate basal radial glial cell (bRGC) expansion, as is seen in the developing human cortex in vivo, and promote astrogliogenesis (Watanabe et al., 2017). At W8, 4G-associated organoids exhibited prominent layered organization with a well-defined ventricular zone (VZ) containing LHX2^+^ PAX6^+^ apical radial glial cells (aRGCs), subventricular zone (SVZ) containing TBR2^+^ intermediate progenitors (IP), and subplate/cortical plate compartment with many CTIP2^+^ neurons (Figure 5E). The layered organization was also reproducible across individual organoids within the group (Figure 5E). By W12, a prominent outer SVZ was apparent with abundant formation of bRGCs distinguished by dispersed cells expressing SOX2, PAX6, HOPX2, and pVIM yet negative for the IP marker TBR2 (Figure 5F). In our previous publication, LIF activation of STAT3 signaling also improved upper-layer neuronal production and laminar organization of the organoids (Watanabe et al., 2017). At W14 and later time points, cerebral organoids derived from the 4G methods similarly showed abundant formation of SATB2^+^ upper layer neurons that were localized outward in the CP (Figure 5G) reminiscent of the distribution of deep and upper layer neurons seen in the gestational week 15.5 human fetal CP (Watanabe et al., 2017). GFAP^+^ astrocytes with characteristic star-shaped morphologies, along with a smaller number of HepaCAM^+^ cells, were also produced (Figure 5H). Together, these experiments illustrate how the addition of a mixture of diverse TGFβ superfamily molecules to undifferentiated n-hPSCs can alter their developmental trajectory and allow the reproducible formation of high-quality cortical organoids that exhibit the cytoarchitectural features of the developing human fetal cortex in vivo.

### The 4G hPSC enhancement method enables efficient production of cerebral, ganglionic eminence, and hippocampal organoids

We lastly tested if 4G-primed hPSCs are biased towards forming cortical organoid structures or broadly capable of forming different telencephalic regions. During development in vivo, organizers secrete patterning molecules to form signaling gradients that, in turn, specify positional identity (Monuki, 2007). For example, in the telencephalon, high levels of SHH provide ventral patterning to form the ganglionic eminences (GE), whereas high concentrations of BMPs/WNTs provide dorsal patterning to form the dorsomedial telencephalon (DMT), which includes the choroid plexus, cortical hem, and hippocampal primordium. Neuroepithelial cells that are not exposed to any of these signals default to forming cerebral cortex. To mimic the ventralizing actions of SHH in vivo, we exposed organoids to varying concentrations of Smoothened Agonist (SAG) at W2.5 (Figure 6A) as previously described (Kadoshima et al., 2013; Watanabe et al., 2017) and compared to untreated samples. In the absence of SAG addition, W5 organoids highly expressed many cortical markers, including FOXG1, NCAD, LHX2, CTIP2, and TBR1 (Figure 6B-6C). Following the addition of 1 μM of SAG, GE markers such as FOXG1, NKX2.1, OLIG2, GSX2, and CTIP2, were highly expressed and cortex-specific markers (LHX2 and TBR1) were suppressed (Figure 6B-6C). By W10, SAG-treated organoids expressed multiple markers of differentiated interneurons including GAD65, somatostatin (SST), and Calretinin, which contrasted with the untreated organoids which lacked these markers and instead exhibited cortical features (Figures 6B-6E). At W14, the mature inhibitory neuronal marker GABA was expressed throughout these ventralized organoids (Figure 6B). Hence, forebrain organoids derived from 4G-treated hPSCs can be readily differentiated into GE-like organoids upon Shh pathway activation.

**Figure 6.**
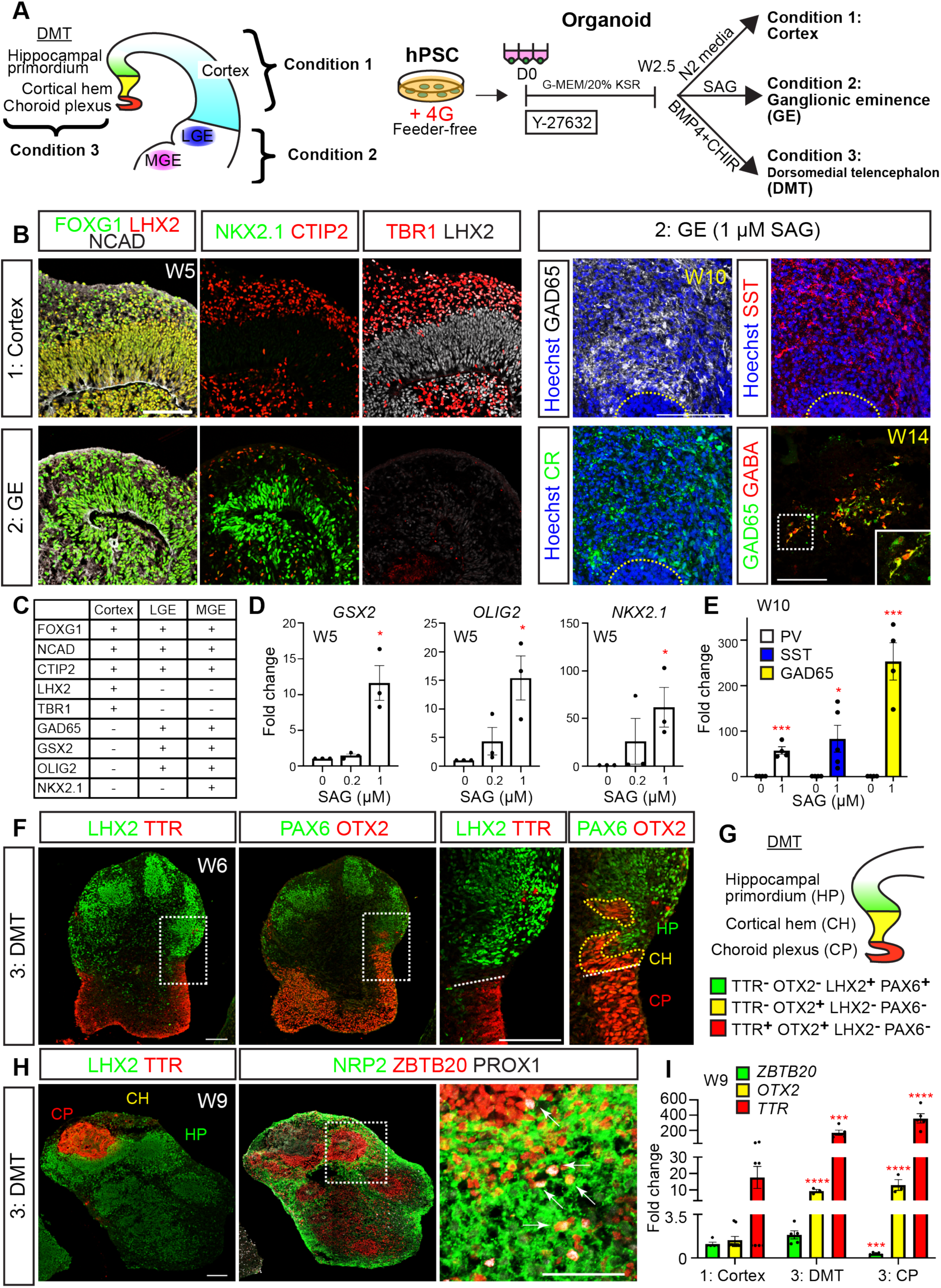
hPSCs cultured under 4G conditions can form multiple telencephalic organoid subtypes: cortex, ganglionic eminence, and dorsomedial telencephalon. (A) Schematics showing a coronal view of the telencephalon and the organoid protocols used to create cortex, ganglionic eminence (GE), and dorsomedial telencephalon (DMT) structures. The DMT includes choroid plexus (CP), cortical hem (CH), and hippocampal primordium (HP). (B) Representative examples of W5 cortical and GE organoids. By default, forebrain organoids exhibited cortical character distinguished by FOXG1, LHX2, NCAD, CTIP2, and TBR1 expression. Organoids exposed to Smoothened Agonist (SAG) instead expressed GE markers such as FOXG1, NKX2.1, and CTIP2 and lacked cortical markers such as LHX2 and TBR1. At W10 and W14, GE organoids expressed many inhibitory neuron markers such as GAD65, somatostatin (SST), calretinin (CR), and GABA. (C) List of cortical, lateral GE (LGE), and medial GE (MGE) markers. (D-E) RT-qPCR analyses of GE progenitor and inhibitory neuron markers. Expression levels are normalized to organoids without SAG application. Data are represented as mean ± SEM. n ≥ 3 independent experimental replicates. (F) DMT induction with the application of BMP4 and CHIR. CP regions are distinguished by TTR and OTX2 staining and form adjacent to CH regions demarcated by OTX2 staining in the absence of TTR. HP, distinguished by LHX2 and PAX6 costaining also formed adjacent to the CH region. (G) Schematic of gene expression patterns seen in the DMT in vivo. (H) HP induction was observed with the application of BMP4 and CHIR. The HP, located next to the cortical hem region, was positive for hippocampal markers such as ZBTB20 and NRP2. Expression of PROX1, a dentate gyrus granule cell marker was also observed. (I) RT-qPCR analyses of DMT markers. Expression levels are normalized to organoids without BMP4 and CHIR application. Data are represented as mean ± SEM. n ≥ 3 independent experimental replicates. All scale bars: 100 μm.

We next dorsalized early forebrain organoids through a combination of BMP and WNT pathway stimulation at W2.5 (6.5 ng/ml BMP4 and 3 μM CHIR), as previously described (Sakaguchi et al., 2015). At W6, we observed formation of distinct dorsomedial telencephalic (DMT) compartments within the organoids including choroid plexus-like regions positive for TTR and OTX2 adjacent to a cortical hem-like region negative for TTR and positive for OTX2 (Figure 6F-6G). Next to these hem-like cells were cells with the characteristics of hippocampal primordium including expression of PAX6, LEF1, NEUROPILIN2 (NRP2), and ZBTB20 (Figure S7A). By W9, expression of ZBTB20 and NRP2 increased and additional hippocampal markers such as the dentate gyrus (DG) granule cell marker PROX1 became evident adjacent to the choroid plexus and cortical hem-like regions (Figures 6H, S7B-S7C, and S7E-S7F). With additional time, expression of the Kainate receptor GLUK1, a characteristic feature of the CA3 region in vivo, also became evident in these organoids (Figures S7F-S7G), which also included choroid plexus and cortical hem-like regions (Figures S7D-S7E). Interestingly, the Kainate receptor GLUK1 positive cells were segregated from PROX1 positive cells, reminiscent of the regional formations of DG and CA3 in vivo (Figures S7E-S7G). Compared to untreated samples, BMP4 and CHIR treated organoids, as well as choroid plexus-like structures that were isolated from these organoids, showed upregulation of DMT markers including OTX2, TTR, LMX1A, and MSX1 (Figures 6I and S7H) by RT-qPCR analyses. ZBTB20 and LHX2 are both expressed in the neocortex and hippocampal primordium but not in the choroid plexus in vivo (Mangale et al., 2008; Sakaguchi et al., 2015). Consistent with in vivo expression patterns, we observed reduced expression of ZBTB20 and LHX2 in the isolated choroid plexus-like structures compared to untreated samples and intact BMP4 and CHIR treated organoids (Figures 6I and S7H). Together, these results demonstrate that n-hPSCs pre-treated with the 4G mixture of TGFβ related growth factors possess broad developmental competence and can be directed to form diverse forebrain structures suitable for disease modeling.

## DISCUSSION

These studies demonstrate, for the first time, that the state of hPSC pluripotency is a significant variable that needs to be considered when conducting brain organoid studies. Pluripotency is a metastable state in vitro and has been primarily described in the context of three major stages: naïve, formative/intermediate, and primed, referring to the pre-, early post-, and later post-implantation epiblasts in vivo. Here, we show that hPSCs maintained under different culture conditions exhibit marked differences in their transcriptional states and developmental potential, spanning the spectrum of naïve and primed pluripotency. Remarkably, we found that the best organoids form from cells that exhibit a particular intermediate state defined by the expression of certain genes associated with TGFβ superfamily signaling and naïve pluripotency. These studies also provide a means for modulating pluripotent states to ensure the overall quality and consistency of organoids across experiments. We further observed that a subset of genes associated with naïve pluripotency, exemplified by *TFAP2C* and *KLF5*, can serve as predictive indicators for efficient organoid formation. Activation of TGFβ superfamily signaling in n-hPSCs can alter their transcriptional profile and boost their capacity to form high-quality organoids.

What is the most effective strategy to ensure good organoid differentiation? Our studies suggest that a critical starting point is to determine the state of pluripotency of a given hPSC population. At one extreme, naïve hPSCs are locked into a program of self-renewal and genomic hypomethylation from which the cells need to be released in order to respond to inductive cues. At the opposite extreme, primed hPSCs are associated with the expression of lineage specific fate determinants that limit the range of cell types into which they can differentiate (Weinberger et al., 2016; Morgani et al., 2017). Indeed, we found that naïve hPSCs were unable to directly form forebrain organoids, consistent with recent studies showing that these cells must first pass through formative/intermediate states in order to undergo lineage-specific differentiation (Figure S1 and Rostovskaya et al., 2019). These formative states appear to be transcriptionally homogeneous and distinct from both naïve and primed states (Kalkan et al., 2017; Smith, 2017; Rostovskaya et al., 2019). Moreover, cell lineages that are established early in embryonic development, such as germline stem cells, are more effectively produced when starting from the formative intermediate state rather than the primed state. Interestingly, key transcription factors expressed during the formative state including OTX2, OCT6, SOX2, and SOX3, are also highly expressed in the early neuroectoderm (Morgani et al., 2017; Smith, 2017), thus suggesting a predisposition for these cells to form neuroectoderm. Perhaps not surprisingly, the cells in the formative state might be prone to neuroectodermal lineage commitment in the absence of BMP/ TGFβ signaling, consistent with the neural default model (De Robertis and Sasai, 1996; Munoz-Sanjuan and Brivanlou, 2002).

We found that hPSCs under our MEF-supported and feeder-free culture conditions are both transcriptionally intermediate, yet they are markedly distinct from each other in their transcriptional states. On a global transcriptomic level, MEF-supported cells appear to be shifted more towards, yet distinct from, naïve hPSCs compared to feeder-free cultures (Figure 3E). Indeed, the feeder-supported cells that we classified as c-hPSCs most closely resembled formative intermediate hPSCs midway through their capacitation trajectory (Figure 3E). These results suggest that there might be particular intermediate hPSC states within the spectrum of naïve and primed pluripotency, some of which are more or less competent to effectively form neuroectodermal organoids. These particular intermediate hPSC states could be neuroectoderm-specific or universal to other lineages. It then follows that as cells become primed, they may become increasingly heterogeneous with subpopulations biased or committed to specific lineage identities including mesoderm and endoderm (Mohammed et al., 2017; Nguyen et al., 2018), which are less favorable for brain organoid formation.

While evidence from our study has demonstrated that it is critical to understand how exogenous factors influence hPSC states, genetic background also matters. Many of the growth factors that influence pluripotency and developmental trajectory are produced by the hPSCs themselves and could vary as a result of intrinsic differences. If the specific genetic background of a given hPSC line were to favor the maintenance of an intermediate state that is more or less permissive to efficient neuroectodermal differentiation, it could potentially influence organoid formation and thereby cloud analysis of how a given disease-associated mutation affects neural development and function. It is therefore imperative to characterize the intrinsic states of hPSC pluripotency across genetically diverse hPSCs and to achieve culture conditions that uniformly maintain organoid competence.

Many patient-derived induced PSCs have been derived under feeder-free conditions due to its convenience and a desire to eliminate exposure of hPSCs to animal products to enable their use in clinical applications (Amit and Itskovitz-Eldor, 2006; Seifinejad et al., 2010). While MEF support remains our benchmark standard for most hPSCs, similar organoid results may be achieved with feeder-free cells by pre-conditioning the cultures with the 4G mixture of TGFβ superfamily growth factors defined in our studies. Furthermore, the 4G method is highly flexible and can be used to form different regions of the forebrain: cortex, ganglionic eminence, and dorsomedial telencephalon including the choroid plexus, cortical hem, and hippocampus.

Our studies further implicate the importance of quality control for the key reagents that can significantly influence the transcriptomic signatures of hPSCs namely MEFs and KSR. When we conducted experiments using different lot numbers of these components, we sometimes observed reduced organoid quality (Figure 1, compare MEFa to MEFb conditions). MEFs secrete many factors, including ACTIVIN A (Figure S2D), which our studies show are both necessary and sufficient to enhance organoid competency when combined with other TGFβ superfamily growth factors. Depending on the MEF batch, the concentrations of ACTIVIN A and other MEF-produced factors might differ and thus influence both hPSC state and forebrain organoid competency. Another key reagent is KSR, which contains two key bioactive ingredients of note: insulin (∼6.7 μg/ml in 10% KSR) and lipid-rich albumin partially purified from bovine serum (AlbuMAX) (Garcia-Gonzalo and Izpisua Belmonte, 2008; Wataya et al., 2008). One out of seven KSR tested batches (∼14%) was optimal for producing forebrain organoids (data not shown), which was consistent with the previously reported mouse cerebral organoid formation (Nasu et al., 2012). Insulin has been shown to caudalize the early neuroectoderm and induce formation of the diencephalon, midbrain, and rostral hindbrain during organoid differentiation (Wataya et al., 2008; Muguruma et al., 2015; Shiraishi et al., 2017). However, insulin’s roles in hPSC transcriptomic signatures have not yet been explored.

By contrast, the roles of cellular lipids and AlbuMAX in hPSCs have been well described. Some studies have demonstrated that AlbuMAX and cellular lipids, such as prostaglandin E2, linoleic acid, and fatty acid synthesis are essential for stem cell pluripotency (Garcia-Gonzalo and Izpisua Belmonte, 2008; Kim et al., 2009; Wang et al., 2017; Dahan et al., 2019). Moreover, a recent report has suggested that lipid deprivation in E8 based-cell culture media could promote a naïve-to-primed intermediate state via changes in the epigenetic landscape, naïve protein expression levels, and mitochondrial morphology (Cornacchia et al., 2019). We observed that hPSCs cultured under feeder-free mTeSR1 culture conditions occupy a similar intermediate transcriptional state as these E8-maintained cells (Figure 3E). When compared to other hPSC populations spanning the spectrum of naïve and primed pluripotency, these and other feeder-free hPSCs appear to be slightly biased towards the primed state compared to feeder-supported intermediate cells which tilt more towards the naïve state of pluripotency (Figure 3E).

Although the transcriptional state of hPSCs can be profoundly impacted by growth under feeder-free or MEF dependent conditions, other contributing factors should not be overlooked. There can still be batch variabilities in other cell culture components as well as procedural differences in hPSC culture techniques. Based on published datasets, we found that MEF-supported H9 ESCs reported by the Smith lab clustered more towards primed hPSCs compared to MEF-supported H9 grown in either the Studer lab or our own lab (Figure 3E). These observations demonstrate how the same hPSC line maintained in different labs can show significant variability in their state of pluripotency, which could, in turn, influence performance in organoid differentiation protocols. The systematic implementation of uniform hPSC maintenance conditions across laboratories may thus be an essential prerequisite to achieve consistent organoid differentiation outcomes.

## EXPERIMENTAL PROCEDURES

### hPSC maintenance and telencephalic organoid differentiation

hPSC experiments were conducted with prior approval from the University of California Los Angeles (UCLA). H9 hESCs and XFiPSCs, were obtained from the UCLA Broad Stem Cell Research Center Core. UCLA1 hESCs harboring homozygous mutation of *TFAP2C* and a dox-inducible TFAP2C expression cassette were used as previously described (Chen et al., 2018; Pastor et al., 2018). hiPSC lines E9 (WT control) and *CHD2* heterozygous indel mutant (Tidball et al., 2017) were obtained by Dr. Jack Parent at the University of Michigan. hPSCs under MEF-supported conditions (Millipore, PMEF-CF) were maintained as previously described (Watanabe et al., 2017). hPSCs under feeder-free conditions were maintained with mTeSR^TM^1 (Stemcell Technologies, 85850) or Essential 8 medium (E8, ThermoFisher, A15117001) and where specified on hESC-qualified Matrigel substrate (Fisher Scientific, 08-774-552). Every 3-5 days, hPSCs were passaged at a 1:10-1:15 dilution with partial dissociation using ReLeSR (Stemcell Technologies, 5872). For the 4G method, hPSCs were preconditioned with growth factors for 3-4 days one day after passage. Growth factor concentrations were as follows: BMP4 (0.1 ng/ml, Invitrogen, PHC9534), TGFβ1 (0.1 ng/ml, R&D Systems, 240-B), ACTIVIN A (10-20 ng/ml, Peprotech, 120-14P), and NODAL (50 ng/ml, R&D Systems, 3218-ND) or TGFβ3 (1 ng/ml, R&D Systems, 8420-B3).

Cerebral and basal ganglionic eminence (GE) organoid formation was performed as described in Watanabe et al., 2017. Of note, KnockOut Serum Replacement (KSR) is known to affect differentiation efficiency and we carried out the differentiation protocol using the same KSR lot (KSR, Invitrogen, 10828010 and 10828028) lot (1670543) as in Watanabe *et al*., 2017. For the formation of dorsomedial telencephalic (DMT) organoids, we used a modified version of previously described methods (Sakaguchi et al., 2015). From day 18-24, BMP4 (6.5 ng/ml, Invitrogen, P2026055) and GSK3 inhibitor (3 μM of CHIR 99021, Fisher Scientific, 442310) were added with DMEM/F-12 (Hyclone) supplemented with 1 % N2 (Invitrogen), 1 % chemically defined lipid concentrate (CDLC, Invitrogen), 10 % fetal bovine serum (FBS qualified source US region, Invitrogen, 10437028) under 40 % O_2_ and 5 % CO_2_ conditions at 37°C. On day 42, DMT organoids were cut in half and the base media was changed to Neurobasal medium (Invitrogen), supplemented with 1 % N2 (Invitrogen), 1 % CDLC, 10 % FBS, 2 % B-27 supplement without vitamin A (Invitrogen), GlutaMAX (Invitrogen), 100 μg/ml of Primocin (InvivoGen). After day 42, DMT organoids were cut into half every two weeks. From day 56, DMT organoids were cultured in Lumox dishes (SARSTEDT). Media was subsequently changed every 2-3 days until the organoids were collected for analysis.

### Primed hPSC maintenance, primed to naïve conversion, and naïve hPSC maintenance

Primed hPSCs were maintained in primed media consisting of 20% KSR in DMEM/F12 supplemented with 1x nonessential amino acids, 2 mM L-Glutamine, 0.5x Penicillin/Streptomycin (all from Invitrogen), 0.1 mM β-mercaptoethanol (Sigma-Aldrich), and 4 ng/ml FGF2 (Peprotech). Primed hPSCs were cultured on CF-1 irradiated MEFs and passaged every 4-5 days with collagenase IV (ThermoFisher Scientific) at 37°C for 5 minutes, followed by manual dissociation by pipetting.

Primed hPSCs were converted to naïve H9 as previously described (Guo et al., 2017). Briefly, primed hPSCs were dissociated into single cells with Accutase and 2×10^5^ cells per 6-well were plated in primed media with 10 μM Y-27632 onto MEFs seeded at a density of 2×10^6^ cells per 6-well plate. The following day (day 1), media was changed to cRM-1, which consists of N2B27 basal media supplemented with 1 μM PD0325901 (Cell Guidance Systems), 20 ng/ml human LIF (Millipore) and 1 mM Valproic Acid (Sigma-Aldrich). On day 4, the media was switched to cRM-2 consisting of N2B27 basal media supplemented with 1 μM PD0325901 (Cell Guidance Systems), 20 ng/ml human LIF (Millipore), 2 μM Gö6983 (Tocris) and 2 μM XAV939. From day 11 onwards, converted naïve cells were cultured in t2iLGö media. Cells were passaged on day 5, day 10, and every 4-5 days subsequently. Homogenous naïve hPSC lines were obtained after 4 passages in t2iLGö media.

Naïve hPSCs were subsequently maintained in t2iLGö media (Takashima et al., 2014) consisting of a 1:1 mixture of DMEM/F12 and Neurobasal, 0.5 % N2 supplement, 1 % B27 supplement, 1x nonessential amino acids, 2 mM L-Glutamine, 0.5x Penicillin/Streptomycin (all from ThermoFisher Scientific), 0.1 mM β-mercaptoethanol (Sigma-Aldrich) (N2B27 basal media) supplemented with 1 μM PD0325901 (Cell Guidance Systems), 1 μM CHIR99021 (Cell Guidance Systems), 20 ng/ml human LIF (Millipore) and 2 μM Gö6983 (Tocris) on CF-1 irradiated MEFs. Naïve hPSCs were passaged every 4 days with Accutase (ThermoFisher Scientific) at 37° C for 5 minutes. Naïve hPSCs between passages 5-7 were used for organoid experiments.

All H9 ESCs were cultured in 5% O_2_, 5% CO_2_ at 37°C and subject to daily media changes.

### Tissue processing and immunohistochemistry

Brain organoids were fixed, cryoprotected, embedded, frozen, and cryosectioned as previously described (Watanabe et al., 2017). Sectioned tissues were collected onto Superfrost Plus slides (Fisher Scientific) and blocked for 30 minutes in PBS with 1 % heat inactivated equine serum (Hyclone), 0.1 % Triton X-100, and 0.05% sodium azide and incubated in primary antibodies (see Table S6) in the blocking solution overnight at 4°C. After three washes in PBST (0.1 % Triton X-100), tissue was incubated with secondary antibodies for one hour at room temperature. After three washes, tissue was mounted in ProLong® Diamond (Invitrogen) with coverslips and stored in the dark at 4°C prior to imaging.

### Microscopic imaging

Confocal images were acquired using a Zeiss LSM 780 or 800 microscope equipped with a motorized stage and Zen black or blue software. Tiled images were assembled using the Zen Tiles with the multi-focus function. For brightfield imaging, a Zeiss Axio Observer D1 microscope was used. All images were compiled in Adobe Photoshop or ImageJ, with image adjustment applied to the entire image and restricted to brightness, contrast, and levels.

### RNA isolation, processing, and RNA-sequencing analyses

Samples were lysed in QIAzol and RNA extracted following manufacturer’s instructions (Qiagen, miRNeasy Micro Kit). For RNA-sequencing analyses, hPSCs, particularly H9 hESCs and XFiPSCs, were collected with 3 replicates (3 independent experiments, 4 conditions, 12 samples total) for feeder-dependent and feeder-independent conditions. RNA integrity was confirmed with the Agilent 2200 TapeStation (RIN > 8) and sent to the UCLA Neuroscience Genomic Core. cDNA libraries were generated using TruSeq with Ribo-Zero Gold (Illumina) and sequenced using an Illumina HiSeq 4000 system, yielding about 50 million reads per sample. We utilized paired end RNA-sequencing with 75 bp reads.

STAR (version 2.4.0j; (Dobin et al., 2013)) was used to align RNA reads to the human genome (GRCh37/hg19). All samples had greater than 80% read alignment. This genome version was also used for subsequent read quantification (exon counts) with HTSeq (version 0.6.1p1; (Anders et al., 2015)). QC statistics were obtained per sample using Picard tools (http://broadinstitute.github.io/picard) to be used in downstream analysis to account for technical variation in gene expression data. QC statistics collected included: AT/GC dropout, read duplication rate, GC bias, read depth, percentage of different genomic regions covered (exons, introns, UTRs, etc.), 5’ end sequencing bias etc.

Differential gene expression was performed in R (https://www.r-project.org/) as follows using HTSeq exon and lncRNA counts. First, genes were filtered such that only genes with a count greater than 10 in at least 80% of samples were retained. The DESeq2 package (Love et al., 2014) was then used to both obtain normalized gene expression data (using varianceStabilizingTransform) and to calculate differentially expressed genes between conditions of interest. Based on the association of Picard QC statistics and other covariates with top normalized gene expression principal components, we included the covariates of condition (hPSC type and feeder type), RIN, RNA concentration, and the first principal component of the Picard sequencing statistics as linear model covariates during differential gene expression calculation. Groups of differentially expressed genes were identified as either significantly up- or down-regulated between conditions with a false discovery rate (FDR) < 5%.

Groups of differentially expressed genes between conditions were subjected to the following enrichment analyses. Gene ontology analysis was performed using Metascape online software (http://metascape.org) (Tripathi et al., 2015). We also used cell type markers of the naïve and primed states (Sahakyan et al., 2017) to find any enrichment in groups of differentially expressed genes using the pSI R package (Xu et al., 2014). Protein-protein interaction enrichment was established using DAPPLE (Rossin et al., 2011). For Figure S2E, GTEx RNA-seq data (gene median RPKM data version 6) was utilized. The Genotype-Tissue Expression (GTEx) Project was supported by the Common Fund of the Office of the Director of the National Institutes of Health, and by NCI, NHGRI, NHLBI, NIDA, NIMH, and NINDS. The data used for the analyses described in this manuscript were obtained from the GTEx Portal on 10/26/17.

Several additional analyses were conducted with a regressed dataset, in which the effects of the following covariates were removed in a linear model of the normalized gene expression data: RIN, RNA concentration, and the first principal component of the Picard QC statistics. With this approach, we could explore a dataset that only retained the effects of the conditions (hPSC type and feeder type) and the linear model residual. Comparison of different sample conditions was achieved through principal component analysis of this regressed dataset. Next, to visualize how our samples related to different states of cell maturity (naïve, intermediate, and primed), we also combined our data with that of (Takashima et al., 2014; Liu et al., 2017; Cornacchia et al., 2019; Rostovskaya et al., 2019) using ComBat in the sva R package (Leek et al., 2012) and then conducted principal component analysis with this merged dataset. We also used hierarchical clustering to group samples based on the scaled expression of the top 150 genes up-regulated in the feeder condition and the top 150 genes up-regulated in the feeder-free condition and visualized this clustering with a heatmap (Figure 3B). We performed similar clustering analyses and heatmap visualizations with SMAD signaling players, naïve, and primed factors (Figure 3D). Clustering and heatmap visualization was conducted with the NMF (Gaujoux and Seoighe, 2010) and the ggplot2 (Wickham, 2016) packages in R.

### Quantitative PCR

Reverse Transcriptase qPCR (RT-qPCR) was performed as previously described (Watanabe et al., 2017). Briefly, total RNA was extracted using a RNeasy Mini or miRNeasy Mini Kit (Qiagen) and >500 ng of total RNA was used for cDNA synthesis for each sample, using the SuperScript IV First-Strand Synthesis System (Invitrogen). For RT-qPCR reaction, LightCycler 480 SYBR Green I Master Mix and exon-spanning primer pairs listed below were used with synthesized cDNA. All primer pairs were validated for 1.8 amplification efficiency as described (Watanabe et al., 2012). All reactions were performed using a Roche LightCycler 480 real-time PCR system in triplicates, and relative expression levels were determined by normalizing the crossing points to the internal reference gene β-ACTIN. Primers used are as follows: β-ACTIN (amplicon size 180 bp) fw 5’-GATCAAGATCATTGCTCCTCCT-3’ rv 5’-GGGTGTAACGCAACTAAGTCA-3’: BMP4 (amplicon size 130 bp) fw 5’-TGGTCTTGAGTATCCTGAGCG-3’ rv 5’-GCTGAGGTTAAAGAGGAAACGA-3’: DPPA3 (amplicon size 119 bp) fw 5’-TGTTACTCGGCGGAGTTCGTA-3’ rv 5’-CCATCCATTAGACACGCAGAAA-3’: GAD65 (amplicon size 140 bp) fw 5’-TTTTGGTCTTTCGGGTCGGAA-3’ rv 5’-TTCTCGGCGTCTCCGTAGAG-3’: GREM1 (amplicon size 96 bp) fw 5’-CGGAGCGCAAATACCTGAAG-3’ rv 5’-GGTTGATGATGGTGCGACTGT-3’: GSX2 (amplicon size 106 bp) fw 5’-ATGTCGCGCTCCTTCTATGTC-3’ rv 5’-CAAGCGGGATGAAGAAATCCG-3’: ID4 (amplicon size 184 bp) fw 5’-CACGTTATCGACTACATCCTGG-3’ rv 5’-TGTCGCCCTGCTTGTTCAC-3’: KLF2 (amplicon size 137 bp) fw 5’-CTACACCAAGAGTTCGCATCTG-3’ rv 5’-CCGTGTGCTTTCGGTAGTG-3’: KLF4 (amplicon size 170 bp) fw 5’-CCCACATGAAGCGACTTCCC-3’ rv 5’-CAGGTCCAGGAGATCGTTGAA-3’: KLF5 (amplicon size 118 bp) fw 5’-CCTGGTCCAGACAAGATGTGA-3’ rv 5’-GAACTGGTCTACGACTGAGGC-3’: KLF9 (amplicon size 220 bp) fw 5’-GCCGCCTACATGGACTTCG-3’ rv 5’-GGATGGGTCGGTACTTGTTCA-3’: KLF17 (amplicon size 81 bp) fw 5’-CCCCTCAGCAAGAGATGACG-3’ rv 5’-GCCTGGCTACCCTTGGAAT-3’: LHX2 (amplicon size 115 bp) fw 5’-TCGGGACTTGGTTTATCACCT-3’ rv 5’-GTTGAAGTGTGCGGGGTACT-3’: LMX1A (amplicon size 199 bp) fw 5’-TCAGAAGGGTGATGAGTTTGTCC-3’ rv 5’-GGGGCGCTTATGGTCCTTG-3’: MSX1 (amplicon size 51 bp) fw 5’-AGTTCTCCAGCTCGCTCAGC-3’ rv 5’-GGAACCATATCTTCACCTGCGT-3’: NANOG (amplicon size 116 bp) fw 5’-TTTGTGGGCCTGAAGAAAACT-3’ rv 5’-AGGGCTGTCCTGAATAAGCAG-3’: NKX2.1 (amplicon size 68 bp) fw 5’-AGCACACGACTCCGTTCTC-3’ rv 5’-GCCCACTTTCTTGTAGCTTTCC-3’: OCT4 (amplicon size 156 bp) fw 5’-GGAGAAGCTGGAGCAAAAC-3’ rv 5’-ACCTTCCCAAATAGAACCCC-3’: OLIG2 (amplicon size 90) fw 5’-ATAGATCGACGCGACACCAG-3’ rv 5’-ACCCGAAAATCTGGATGCGA-3’: OTX2 (amplicon size 154 bp) fw 5’-AGAGGACGACGTTCACTCG-3’ rv 5’-TCGGGCAAGTTGATTTTCAGT-3’: PV (amplicon size 96 bp) fw 5’-AAGAGTGCGGATGATGTGAAG-3’ rv 5’-GCCTTTTAGGATGAATCCCAGC-3’: SOX2 (amplicon size 155 bp) fw 5’-GCCGAGTGGAAACTTTTGTCG-3’ rv 5’-GGCAGCGTGTACTTATCCTTCT-3’: SST (amplicon size 108 bp) fw 5’-ACCCAACCAGACGGAGAATGA-3’ rv 5’-GCCGGGTTTGAGTTAGCAGA-3’: TFAP2C (amplicon size 114 bp) fw 5’-CTGTTGCTGCACGATCAGACA-3’ rv 5’-CTCAGTGGGGTTCATTACGGC-3’: TFCP2L1 (amplicon size 245 bp) fw 5’-ATACCAGCCGTCCTATGAAACC-3’ rv 5’-ACTGCGAGAACCTGTTGCG-3’: TGFβ1 (amplicon size 209 bp) fw 5’-CTAATGGTGGAAACCCACAACG-3’ rv 5’-TATCGCCAGGAATTGTTGCTG-3’: TGFβ3 (amplicon size 114 bp) fw 5’-ACTTGCACCACCTTGGACTTC-3’ rv 5’-GGTCATCACCGTTGGCTCA-3’: TTR (amplicon size 229 bp) fw 5’-ATCCAAGTGTCCTCTGATGGT-3’ rv 5’-GCCAAGTGCCTTCCAGTAAGA-3’: ZBTB20 (amplicon size 77 bp) fw 5’-GACAGGATCTACTCGGCACTC-3’ rv 5’-ACTGCGCCGCTGTAAAAAGA-3’: and ZEB1 (amplicon size 86 bp) fw 5’-GATGATGAATGCGAGTCAGATGC-3’ rv 5’-ACAGCAGTGTCTTGTTGTTGT-3’.

### Immunoblotting

For the positive and negative controls, UCLA1 hESCs harboring a homozygous *TFAP2C* deletion or a doxycycline-inducible *TFAP2C* expression cassette were used as previously described (Pastor et al., 2018). In all cases, hPSCs were first washed with cold PBS and then collected for protein in RIPA lysis buffer with protease inhibitors (ThermoFisher, 78425) and phosphatase inhibitors (ThermoFisher, 78420). RIPA buffer was added to the cells followed by 5 minutes incubation on ice. Cells were then scraped off from the culture dishes, transferred to a microcentrifuge tube, and rotated for 10 minutes at 4°C. If a lysate was too viscous, it was sonicated. Lysates were centrifuged at maximum speed for 15 minutes at 4°C and the supernatant collected and snap frozen in liquid nitrogen followed by storage at −80°C. Protein lysates were then mixed with loading dye containing β-mercaptoethanol and placed in a 95°C shaking heating block followed by centrifugation at maximum speed for 2 minutes. The heated lysates were run on SurePage gels (GenScript, M00652) in MOP buffer (GenScript, M00138) for 45 minutes to 1 hour at 120 V, and proteins then transferred onto 0.45 μm nitrocellulose membranes in Tris Glycine buffer for 2 hours on ice at 300 mA. Protein transfer membranes were blocked in 5 % skim milk (Bio-Rad, 170-6404) for 1 hour and incubated with primary antibodies (rabbit anti TFAP2C/AP2 Abcam GR59885-7 at 1:1000 and goat anti GAPDH Abcam ab94583 at 1:1000) on a shaker at 4°C overnight. After three 5-minute washes, membranes were incubated with secondary antibodies conjugated with HRP in 5 % skim milk on the shaker for 2 hours at room temperature. We used SuperSignal West Femto Maximum Sensitivity Substrate (ThermoFisher, 34095) or Pierce ECL 2 Western Blotting Substrate (ThermoFisher, 80197) for chemiluminescent detection. Membranes were scanned using a Sapphire RGBNIR^TM^ Biomolecular Imager (Azure Biosystems, Inc.), and acquired digital images quantified using the ImageJ gel function. TFAP2C protein levels were normalized to GAPDH and reported as mean ± SEM for at least four biological replicates.

### Mass spectrometry

Previous work demonstrated that protein factors secreted by MEFs might be responsible for reprogramming hPSC metabolism (Gu et al., 2016). To understand which protein factor(s) in the MEF-conditioned medium might impact the ability of hPSCs to effectively form cortical organoids, a mass spectrometry-based protein identification experiment was conducted as follows: Conditioned media was collected after 24 hours incubation with or without MEFs. The protein fractions of the samples were enriched by using an Amicon Ultra-15 Centrifugal Filter Unit with Ultracel-10 membrane (Millipore, UFC901024). Concentrated protein samples were then separated by SDS-PAGE and the gel was processed for Coomassie Brilliant Blue staining (Thermo Scientific) according to the manufacturer’s instructions. Coomassie Blue-stained bands were cut from the gels, washed twice with 50% acetonitrile, and processed for liquid chromatography tandem mass spectrometry analysis at the mass spectrometry core facility at Beth Israel Deaconess Medical Center (Boston, MA).

## AUTHOR CONTRIBUTIONS

M.W., N.V., F.T., J.E.B., and O.A.M. performed all organoid culture experiments and coordinated on various analytical procedures. J.H., J.E.B., S.S. and M.J.G. performed bioinformatics analyses. W.G. and H.R.C. contributed mass spectrometry analyses of MEF-secreted factors. A.J.C., D.C., K.P., and A.T.C. provided valuable reagents, datasets, and guidance in the naïve pluripotent stem cell experiments and analyses. M.W. and B.G.N. conceived and designed the experiments with helpful input from the other authors. M.W., J.E.B., and B.G.N. wrote the manuscript with input from the other authors.

## ACCESSION NUMBERS

All RNA-sequencing data reported in our study have been deposited within the Gene Expression Omnibus (GEO) repository and will be accessible through the GEO Series accession number GSE140057 (https://www.ncbi.nlm.nih.gov/geo/) after publication. To access GEO accession GSE140057 before publication, please contact the corresponding author for a reviewer’s token key. The proteomic analyses were deposited to PRIDE Archive (https://www.ebi.ac.uk/pride/archive/) with the accession number of PXD013662.

## ACKNOWLEDGEMENTS

We thank Samantha Butler, Jack Parent, Harley Kornblum, Ranmal Samarasinghe and members of Novitch lab for invaluable discussions and comments on the manuscript, and Jack Parent for the gift of hiPSC lines. M.W. is thankful to Hideya Sakaguchi for sharing and offering advice on the hippocampal organoid protocol. This work was supported by the UCLA Comprehensive Cancer Center and Eli and Edythe Broad Center of Regenerative Medicine and Stem Cell Research (BSCRC) Ablon Scholars Program, research awards from the Rose Hills Foundation, the California Institute for Regenerative Medicine (CIRM) (DISC1-08819) and the NIH (R01NS089817 and R01NS085227) to B.G.N. M.W. was supported by postdoctoral training awards provided by the UCLA BSCRC, the Uehara Memorial Foundation, UCLA Brain Research Institute, and award from the NIH (K99HD096105). J.E.B., N.V., and F.T. were supported partly by the UCLA-California State University Northridge CIRM-Bridges training program (TB1-00183), and a BSCRC predoctoral training award to J.E.B. W.G. was supported by a UCLA Dissertation Year Fellowship. A.T.C. was supported by a grant from the NIH (R01HD079546). K.P. was supported by research awards from the UCLA BSCRC, the David Geffen School of Medicine, the UCLA Jonnson Comprehensive Cancer Center, the NIH (R01HD098387), and a Faculty Scholar grant from the Howard Hughes Medical Institute. M.J.G. was supported by a SFARI Bridge to Independence Award. We also acknowledge the support of the NINDS Informatics Center for Neurogenetics and Neurogenomics (P30NS062691) and both Cells, Circuits, and Systems Analysis and Genetics and Genomics Cores of the Semel Institute of Neuroscience at UCLA, which are supported by the NICHD (U54HD087101).

---

**Figure S1 (related to Figures 1 and 2).**
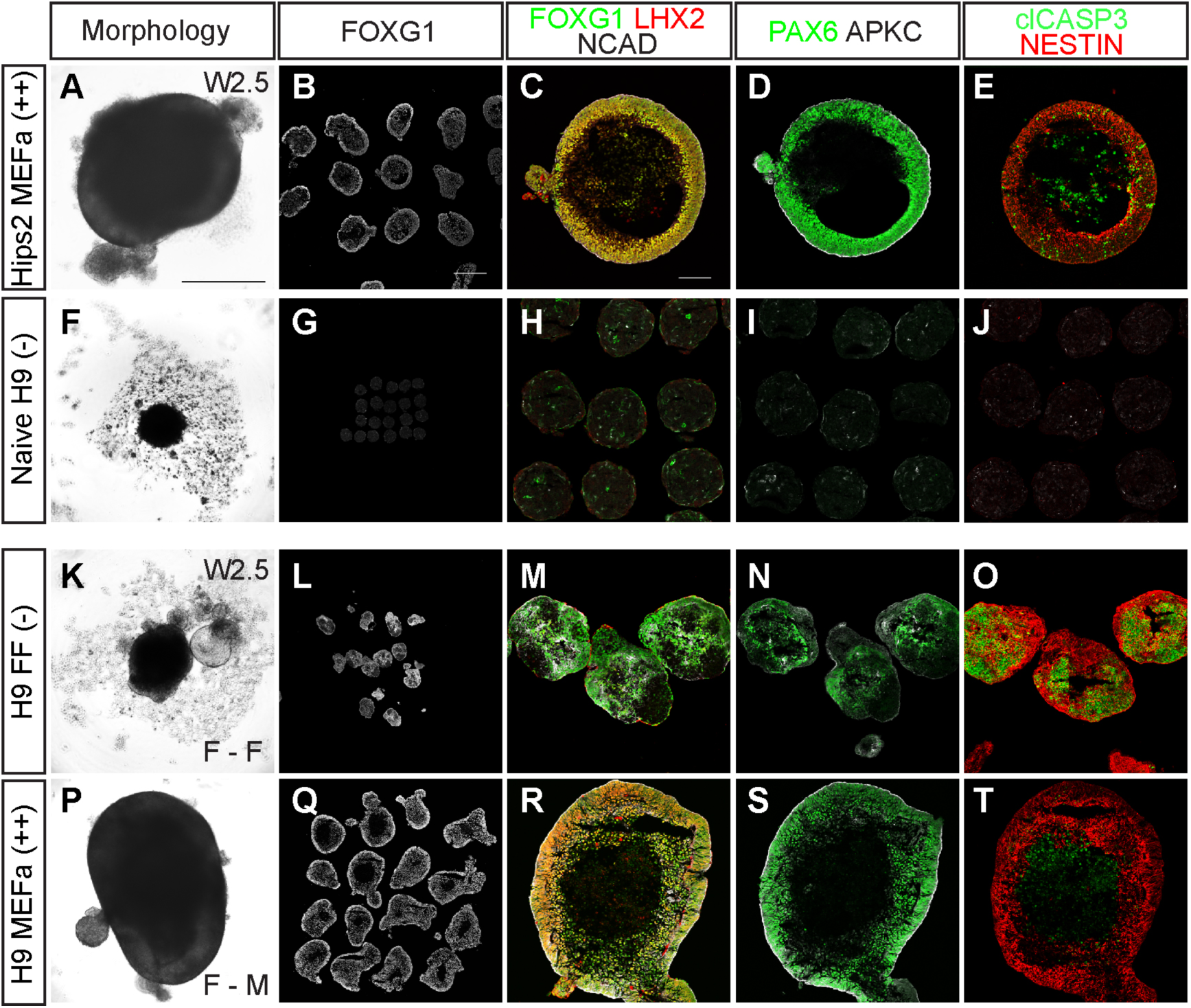
Forebrain organoid differentiation from hPSCs maintained under different cell culture conditions. (A-E) Hips2 hiPSCs cultured under MEFa and KSRa conditions successfully differentiated into forebrain organoids similar to H9 hESCs maintained under the culture conditions shown in Fig. 1C-G. (F-J) H9 naïve cells displayed extremely poor differentiation into forebrain organoids. (K-T) H9 hESCs maintained under feeder-free conditions were adapted to MEFa-dependent conditions. MEFa-adapted H9 efficiently formed forebrain organoids as shown in XFiPSCs in Fig. 2. Scale bars: 500 μm (A, B), 100 μm (C).

**Figure S2 (related to Figure 3).**
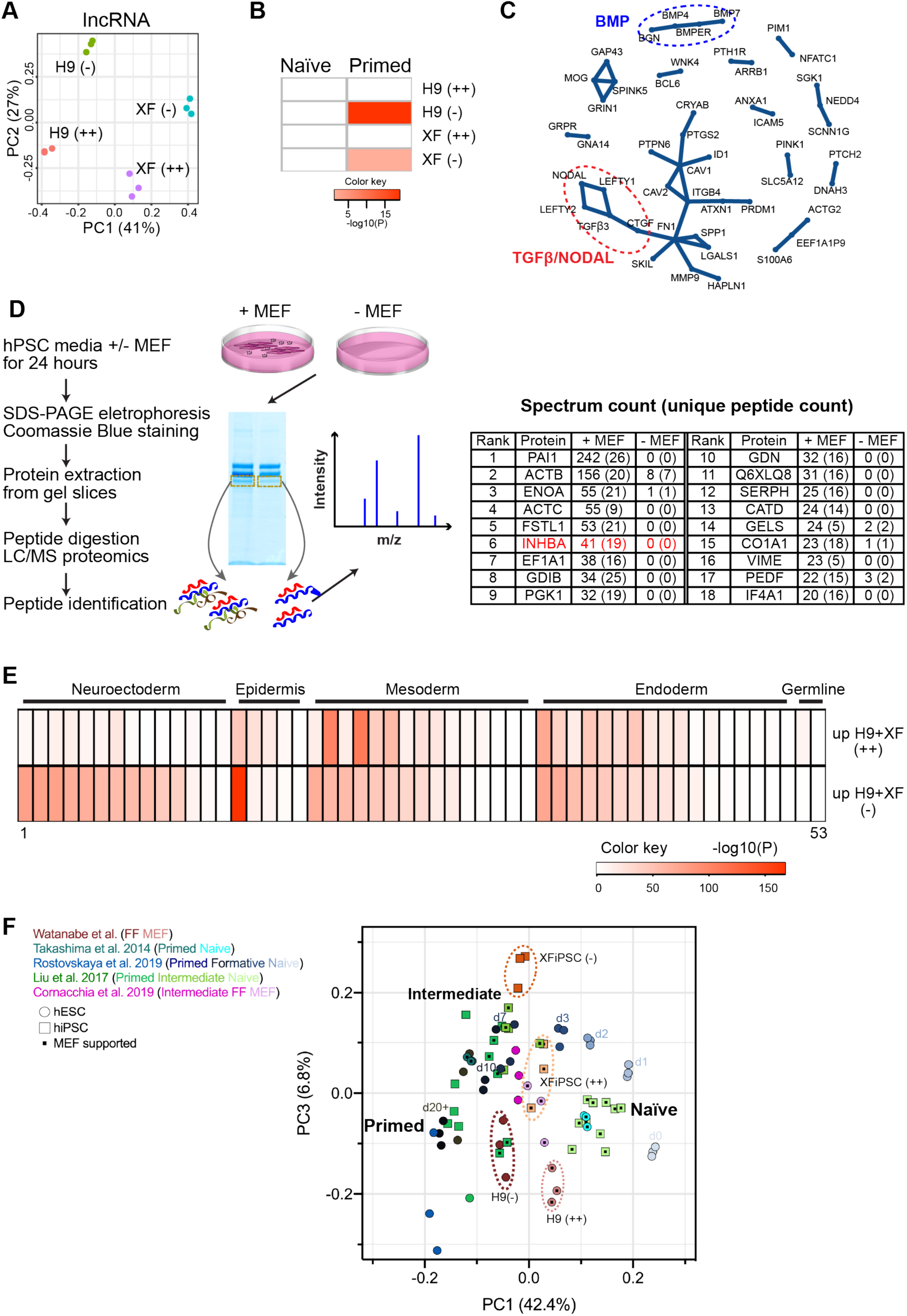
Transcriptomic analysis of hPSCs under different culture conditions and MEF-secreted proteins identified by mass spectrometry. (A) Principal component analysis of lncRNA expressed by H9 hESCs and XFiPSCs maintained under MEFa-dependent conditions associated with effective forebrain organoid formation (++) or feeder-free conditions associated with poor organoid formation (-). Triplicate samples are displayed. (B) Cell Marker enrichment analysis (p<0.001) using genes associated with naïve or primed hPSCs. (C) Protein-protein interaction analysis using Disease Association Protein-Protein Link Evaluator (DAPPLE). (D) Schematic of proteomic analysis and list of MEF-secreted proteins identified by mass spectrometry. (E) Cell marker enrichment analysis (p<0.001) using genes associated with mature cell lineages. A list of the cell lineages displayed (1-53) is provided in Table S5. (F) Principal component analysis integrating previously published data sets with primed, formative/intermediate, and naïve hiPSCs (Takashima et al., 2014; Liu et al., 2017; Cornacchia et al., 2019; Rostovskaya et al., 2019), showing PC1 vs PC3 related to the PC1 vs PC2 plot shown in Fig. 3E.

**Figure S3 (related to Figure 3E).**
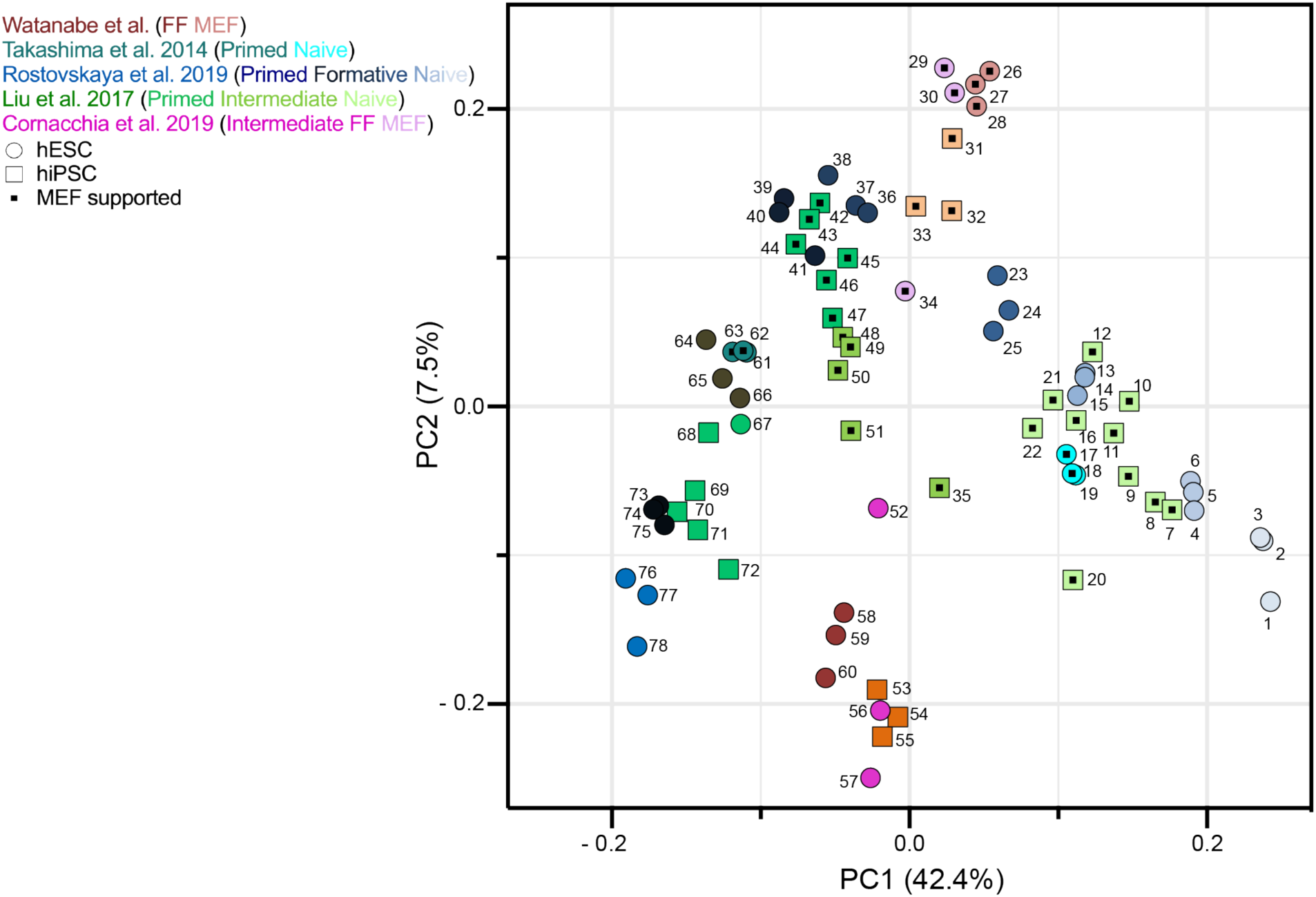
Sample key for principal component analysis shown in Figure 3E. The figure displays each individual data point identified according to the sample listing provided in Table S4. The datasets displayed are described in Takashima et al. 2014, Liu et al. 2014, Cornacchia et al. 2019, and Rostovskaya et al. 2019.

**Figure S4 (related to Figure 3).**
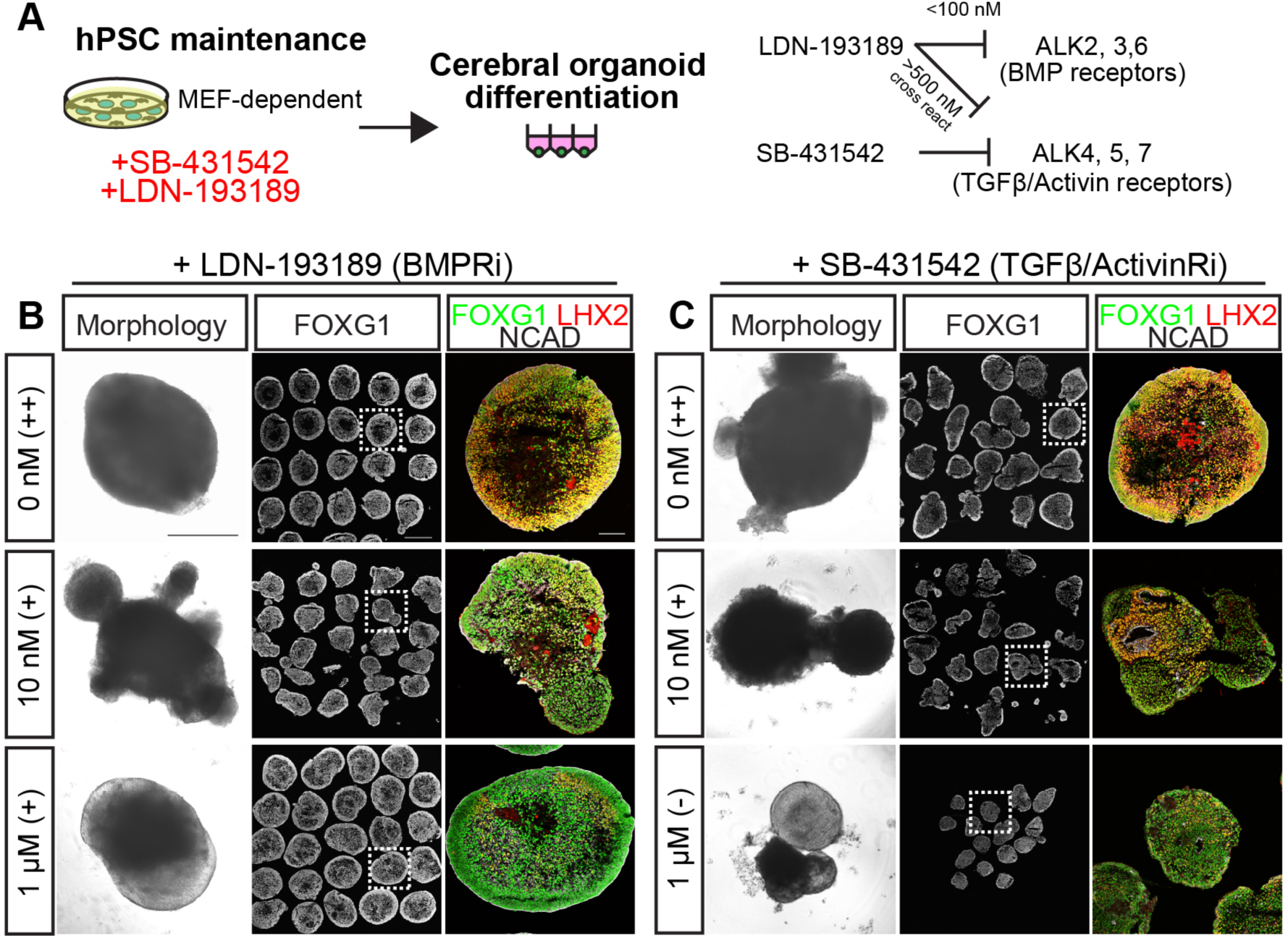
TGFβ signaling in hPSCs is required for optimal forebrain organoid formation. (A) Schematic of the experimental design. LDN-193189 or SB-431542 were added to hPSCs for five days before differentiation. LDN-193189 selectively inhibits BMP receptors when used at concentrations below 100 nM. Above 500 nM, LDN-193189 becomes less selective and also inhibits TGFβ and ACTIVIN receptors (Yu et al., 2008). SB-431542 is a selective inhibitor for TGFβ and ACTIVIN receptors (Inman et al., 2002). Assessment of forebrain organoid formation at W2.5 from MEFa-supported organoid-competent hPSCs with and without SB-431542 (B) or LDN-193189 (C) application. Scale bars: 500 μm, morphology and FOXG1 images; 100 μm FOXG1 LHX2 NCAD images.

**Figure S5 (related to Figures 3, 5, and S2).**
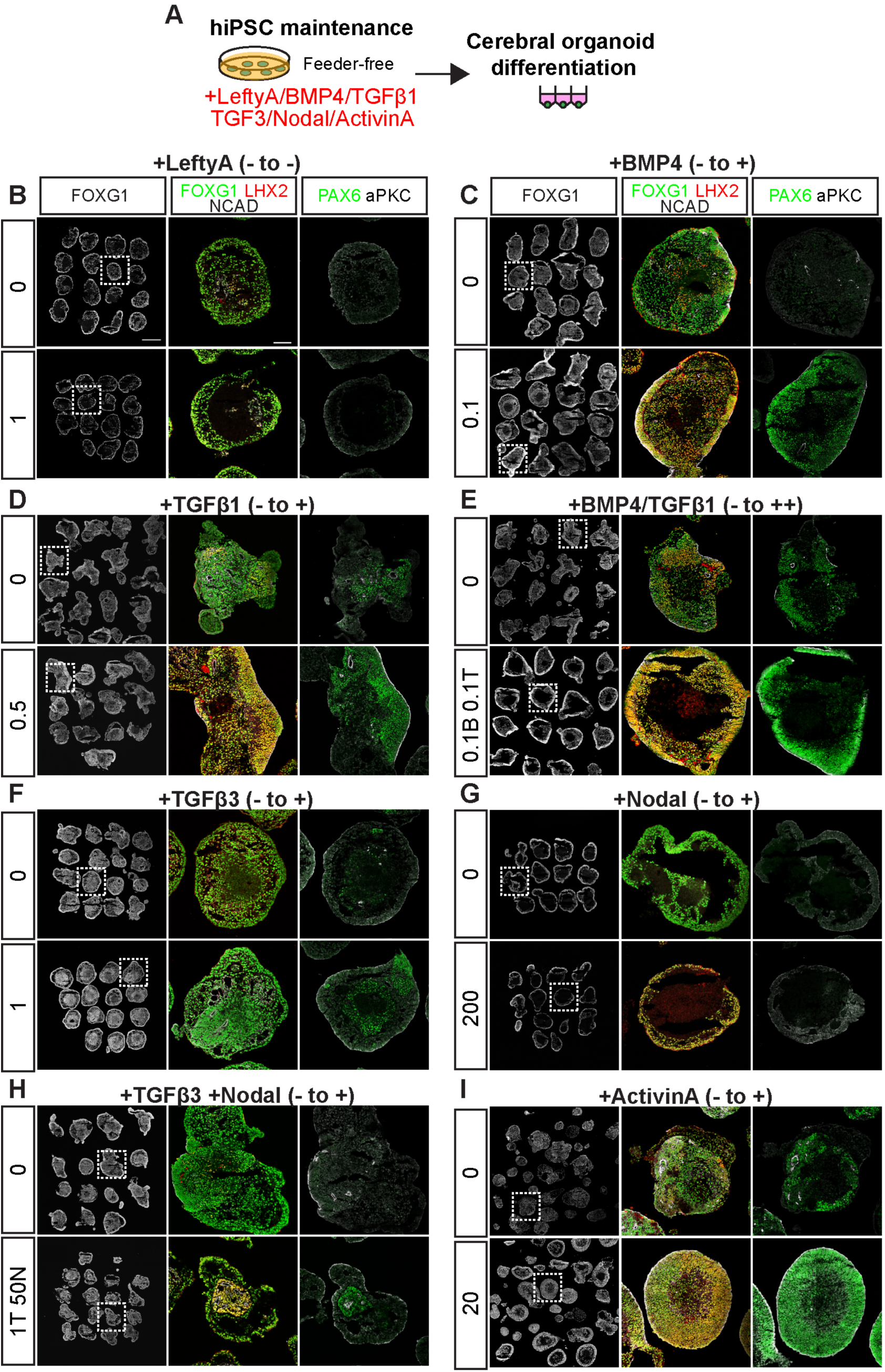
The addition of individual or pairs of TGFβ superfamily growth factors to feeder-free hPSCs partially rescues forebrain organoid formation. (A) Schematic of the experimental design. Organoid non-competent feeder-free H9 were preconditioned for five days with TGFβ superfamily growth factors found to be associated with organoid-competent hPSCs through the transcriptomic and proteomic analyses conducted in Figures 3 and S2. (B-I) Representative examples of W2.5 organoids differentiated from H9 hESC maintained under feeder-free conditions without or with LEFTYA, BMP4, TGFβ1, BMP4/TGFβ1, TGFβ3, NODAL, TGFβ3/NODAL, or ACTIVIN A as indicated. Supplementation with most TGFβ superfamily molecules partially rescued forebrain organoid production with the exception of LEFTYA. All concentrations displayed are in ng/ml. For the average scores and ratings, see also Table S1. Scar bars: 500 μm for FOXG1 images; 100 μm for FOXG1 LHX2 NCAD images.

**Figure S6 (related to Figure 5).**
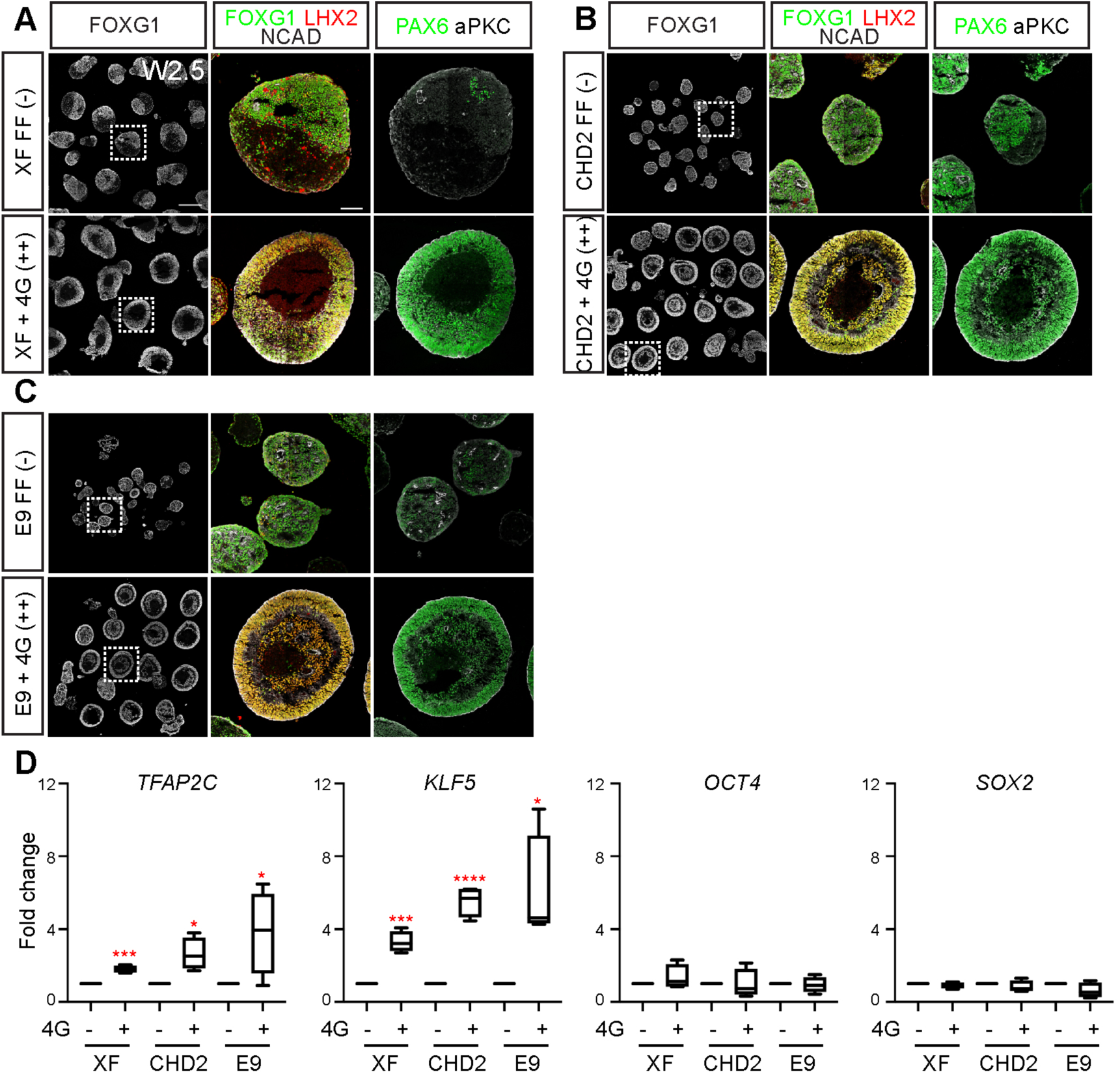
The 4G combination of TGFβ superfamily molecules enhanced forebrain organoid formation from multiple feeder-free hPSC lines. Supplementation with four growth factors (4G)-BMP4, TGFβ1, ACTIVIN A, and TGFβ3 enhanced forebrain organoid formation from feeder-free (A) XFiPSCs, (B) *CHD2* mutant hiPSCs, and (C) E9 healthy patient hiPSCs. (D) RT-qPCR analyses showing the upregulation of *TFAP2C* and *KLF5* with the 4G application. The general pluripotency markers *OCT4* and *SOX2* were unchanged. Expression levels are normalized to feeder-free hiPSCs without 4G addition. Data are represented as mean ± SEM. n ≥ 4 independent experimental replicates. Scale bars: 500 μm FOXG1; 100 μm FOXG1 LHX2 NCAD in (A).

**Figure S7 (related to Figure 6).**
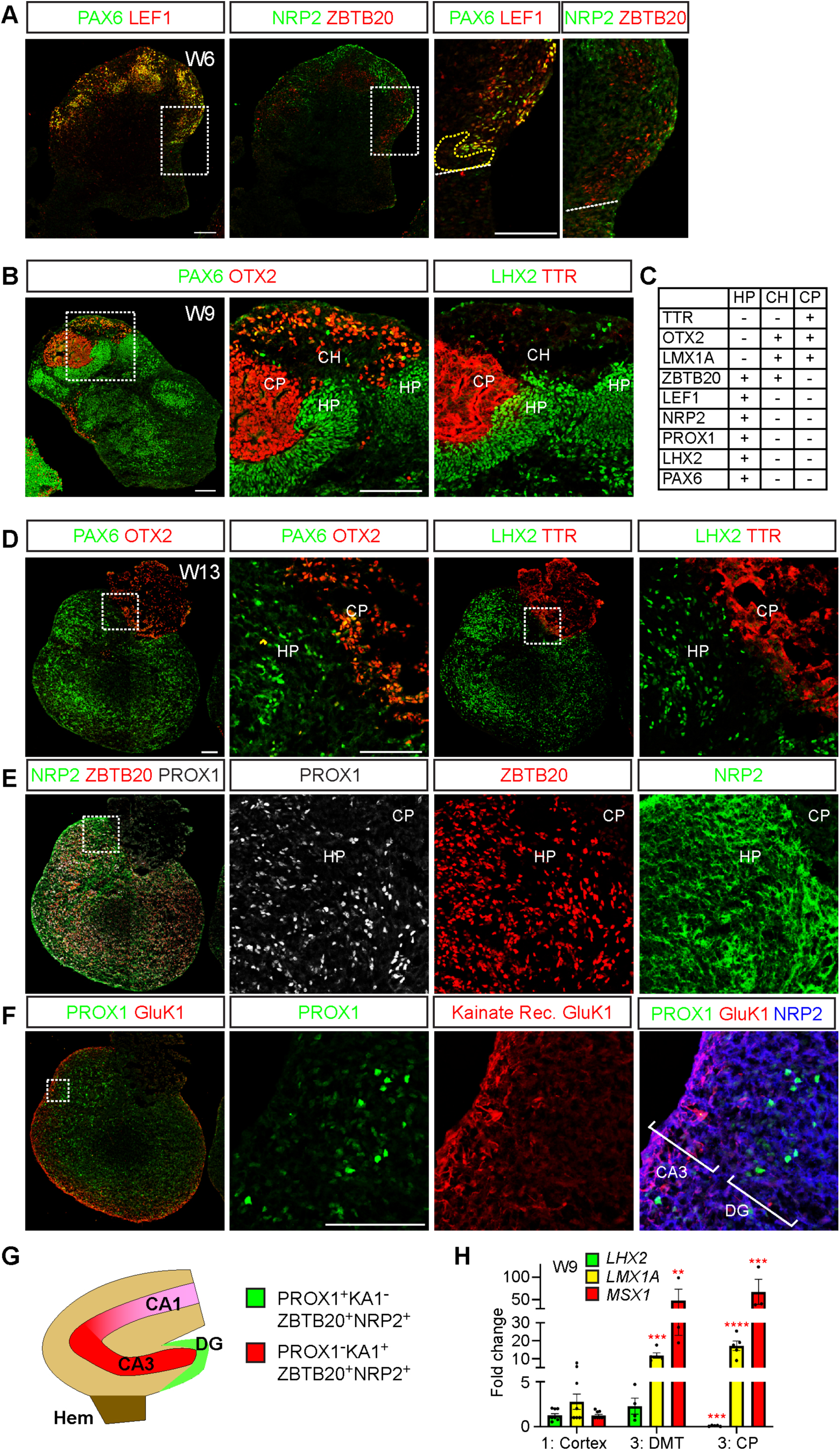
hPSCs cultured under 4G feeder-free conditions can form dorsomedial telencephalic organoids. (A) Representative examples of W6 dorsomedial telencephalic (DMT) organoids stained for hippocampal primordium (HP) markers such as PAX6, LEF1, NRP2, and ZTBT20. Images shown are adjacent sections to Figure 6F. (B) W9 DMT organoids stained for HP (PAX6 and LHX2), cortical hem (CH, OTX2), and choroid plexus (CP, OTX2 and TTR) markers. (C) List of HP, CH, and CP markers. (D-F) Representative examples of W13 DMT organoids. All images are from adjacent sections immunostained for CH (OTX2), CP (OTX2 and TTR), and HP (PAX6, LHX2, NRP2, ZBTB20, PROX1, and Kainate Receptor GluK1) markers. GluK1 staining, which is associated with CA3 axons in vivo, was routinely found adjacent to dentate gyrus (DG)-like regions demarcated by PROX1 expression. (G) Schematic of the organization of the fetal hippocampus in vivo, illustrating the relative positions of the DG, CA3, and CA1 regions. (H) RT-qPCR analyses of DMT markers. Expression levels are normalized to organoids without BMP4 and CHIR application. Data are represented as mean ± SEM. n ≥ 3 independent experimental replicates. All scale bars: 100 μm.

**Table S1.** Average scores and ratings for different culture conditions related to Figures 1, 2, 4, 5, S1, S3, S4, and S5

|  |  |  |  |  |  |  |
| --- | --- | --- | --- | --- | --- | --- |
| Cell line | Culture condition | hESC/<br>hiPSC? | Experimental<br>replicates | Total # of<br>organoids | Avg.<br>score | Avg.<br>rating |
| H9 | MEFa | hESC | 7 | 96 | 8.86 | ++ |
| H9 | MEFb | hESC | 3 | 58 | 6.00 | + |
| H9 | FF | hESC | 17 | 264 | 3.59 | - |
| H9 | Naïve MEFc | hESC | 3 | 78 | 3.00 | - |
| H9 | LEFTYA (1 ng/ml) | hESC | 3 | 39 | 4.00 | - |
| H9 | BMP4 (0.1 ng/ml) | hESC | 3 | 44 | 6.67 | + |
| H9 | TGFβ1 (0.5 ng/ml) | hESC | 3 | 40 | 6.67 | - |
| H9 | BMP4 (0.1 ng/ml)/<br>TGFβ1 (0.1 ng/ml) | hESC | 6 | 85 | 5.83 | + |
| H9 | TGFβ3 (1 ng/ml) | hESC | 3 | 44 | 5.00 | + |
| H9 | Nodal (200 ng/ml) | hESC | 3 | 45 | 5.00 | + |
| H9 | TGFβ1 (1 ng/ml)/<br>Nodal (50 ng/ml) | hESC | 3 | 48 | 6.00 | + |
| H9 | ActivinA (20 ng/ml) | hESC | 3 | 71 | 7.33 | + |
| H9 | FF + #4 (+3G) | hESC | 5 | 91 | 5.80 | + |
| H9 | FF + #7 (+4G) | hESC | 5 | 59 | 8.40 | ++ |
| H9 | FF + #8 (+4G) | hESC | 3 | 35 | 8.00 | ++ |
| H9 | FF + #9 (+5G) | hESC | 4 | 46 | 7.50 | + |
| Hips2 | MEFa | hiPSC | 8 | 96 | 8.25 | ++ |
| XF | FF | hiPSC | 5 | 88 | 3.80 | - |
| XF | MEFa | hiPSC | 12 | 212 | 8.25 | ++ |
| XF | LDN (10 nM) | hiPSC | 3 | 65 | 5.33 | + |
| XF | LDN (1 μM) | hiPSC | 3 | 65 | 7.00 | + |
| XF | SB (10 nM) | hiPSC | 3 | 56 | 5.66 | + |
| XF | SB (1 μM) | hiPSC | 3 | 37 | 3.33 | - |
| XF | E8 | hiPSC | 3 | 46 | 3.33 | - |
| XF | FF + 4G (#7) | hiPSC | 4 | 51 | 8.00 | ++ |
| E9 | FF | hiPSC | 6 | 54 | 3.00 | - |
| E9 | FF + 4G (#7) | hiPSC | 4 | 62 | 8.00 | ++ |
| CHD2 | FF | hiPSC | 7 | 110 | 3.14 | - |
| CHD2 | FF + 4G (#7) | hiPSC | 4 | 61 | 8.25 | ++ |

Tables S2, S3, S4 (see Excel files)

**Table S5.** List of cell types, related to Figure S2C

| Table S5. List of cell types, related to Figure S2C |  |  |  |
| --- | --- | --- | --- |
| # | Description | Origin | Other origin |
| 1 | Brain - amygdala | Neuroectoderm |  |
| 2 | Brain - anterior cingulate cortex | Neuroectoderm |  |
| 3 | Brain - frontal cortex | Neuroectoderm |  |
| 4 | Brain - substantia nigra | Neuroectoderm |  |
| 5 | Brain - hippocampus | Neuroectoderm |  |
| 6 | Brain - putamen/basal ganglia | Neuroectoderm |  |
| 7 | Brain - spinal cord cervical | Neuroectoderm |  |
| 8 | Brain - cortex | Neuroectoderm |  |
| 9 | Brain - hypothalamus | Neuroectoderm |  |
| 10 | Brain - nucleus accumbens/basal ganglia | Neuroectoderm |  |
| 11 | Brain - caudate basal ganglia | Neuroectoderm |  |
| 12 | Brain - cerebellar hemisphere | Neuroectoderm |  |
| 13 | Brain - cerebellum | Neuroectoderm |  |
| 14 | Nerve - tibial | Neural crest | Neuroectoderm |
| 15 | Cells transformed fibroblasts | Epidermis |  |
| 16 | Skin - sun exposed lower leg | Epidermis |  |
| 17 | Skin - not sun exposed suprapubic | Epidermis |  |
| 18 | Breast - mammary tissue | Epidermis |  |
| 19 | Pituitary | Ectoderm |  |
| 20 | Adrenal gland | Mesoderm | Ectoderm |
| 21 | Artery - tibial | Mesoderm |  |
| 22 | Cells EBV transformed lymphocytes | Mesoderm |  |
| 23 | Muscle - skeletal | Mesoderm |  |
| 24 | Artery - aorta | Mesoderm |  |
| 25 | Whole blood | Mesoderm |  |
| 26 | Adipose - visceral/omentum | Mesoderm |  |
| 27 | Artery - coronary | Mesoderm |  |
| 28 | Adipose - subcutaneous | Mesoderm |  |
| 29 | Kidney - cortex | Mesoderm |  |
| 30 | Uterus | Mesoderm |  |
| 31 | Vagina | Mesoderm |  |
| 32 | Cervix - endocervix | Mesoderm |  |
| 33 | Cervix - ectocervix | Mesoderm |  |
| 34 | Fallopian tube | Mesoderm |  |
| 35 | Heart - left ventricle | Endoderm |  |
| 36 | Esophagus - mucosa | Endoderm |  |
| 37 | Liver | Endoderm |  |
| 38 | Esophagus - muscularis | Endoderm |  |
| 39 | Heart - artial appendage | Endoderm |  |
| 40 | Esophagus - gastroesophageal junction | Endoderm |  |
| 41 | Colon - sigmoid | Endoderm |  |
| 42 | Stomach | Endoderm |  |
| 43 | Pancreas | Endoderm |  |
| 44 | Colon - transverse | Endoderm |  |
| 45 | Lung | Endoderm |  |
| 46 | Bladder | Endoderm | Mesoderm |
| 47 | Spleen | Endoderm |  |
| 48 | Minor salivary gland | Endoderm |  |
| 49 | Small intestin - terminal ileum | Endoderm |  |
| 50 | Thyroid | Endoderm | Neural crest |
| 51 | Prostate | Endoderm |  |
| 52 | Ovary | Germline |  |
| 53 | Testis | Germline |  |

**Table S6.**
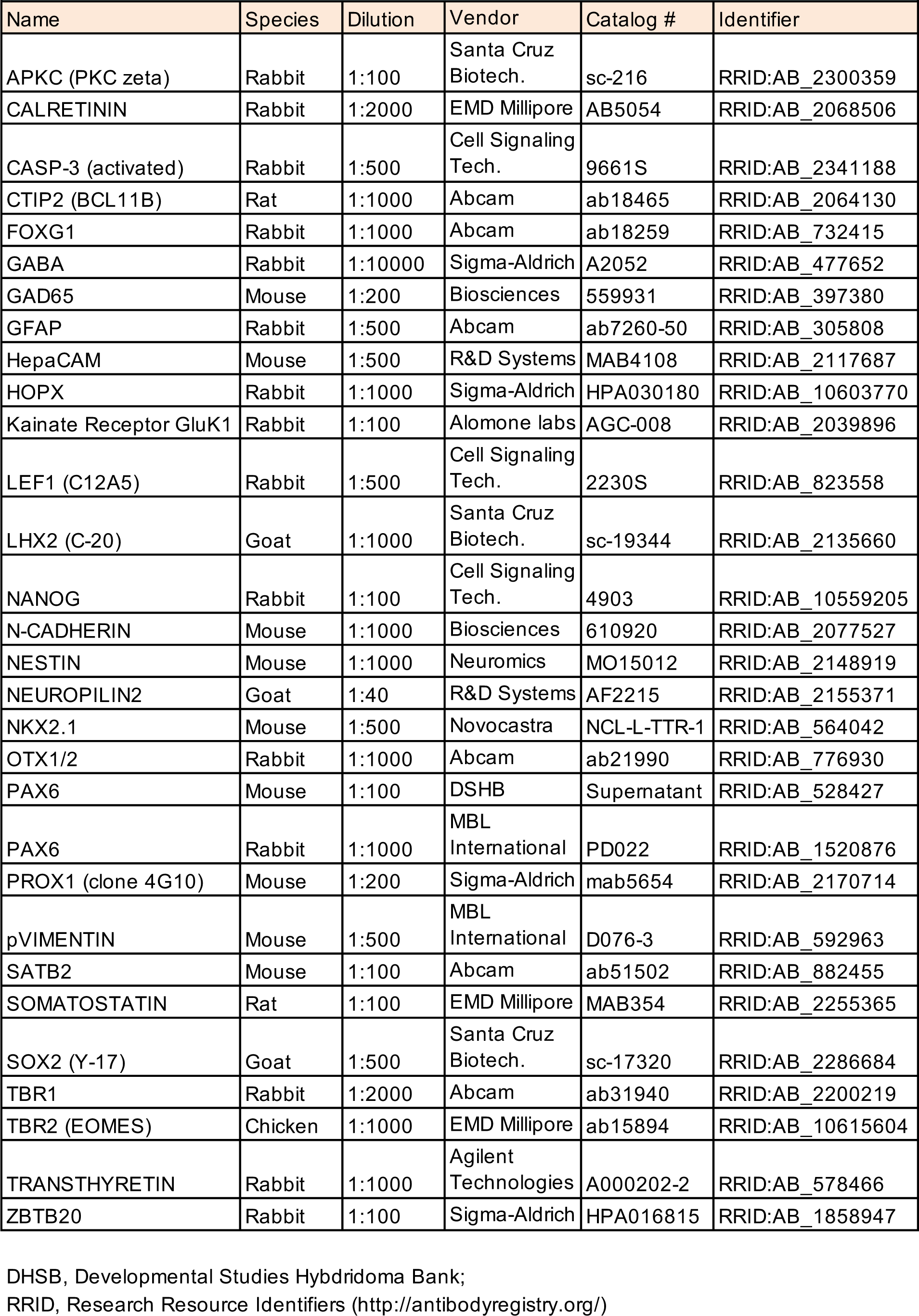
Primary antibody list

## REFERENCES

1. Amit, M., and Itskovitz-Eldor, J. (2006). Feeder-free culture of human embryonic stem cells. Methods Enzymol 420, 37–49.

2. Anders, S., Pyl, P.T., and Huber, W. (2015). HTSeq—a Python framework to work with high-throughput sequencing data. Bioinformatics 31, 166–169.

3. Bagley, J.A., Reumann, D., Bian, S., Levi-Strauss, J., and Knoblich, J.A. (2017). Fused cerebral organoids model interactions between brain regions. Nat Methods 14, 743–751.

4. Bershteyn, M., Nowakowski, T.J., Pollen, A.A., Di Lullo, E., Nene, A., Wynshaw-Boris, A., and Kriegstein, A.R. (2017). Human iPSC-Derived Cerebral Organoids Model Cellular Features of Lissencephaly and Reveal Prolonged Mitosis of Outer Radial Glia. Cell Stem Cell 20, 435–449 e434.

5. Birey, F., Andersen, J., Makinson, C.D., Islam, S., Wei, W., Huber, N., Fan, H.C., Metzler, K.R.C., Panagiotakos, G., Thom, N., et al. (2017). Assembly of functionally integrated human forebrain spheroids. Nature 545, 54–59.

6. Camp, J.G., Badsha, F., Florio, M., Kanton, S., Gerber, T., Wilsch-Brauninger, M., Lewitus, E., Sykes, A., Hevers, W., Lancaster, M., et al. (2015). Human cerebral organoids recapitulate gene expression programs of fetal neocortex development. Proc Natl Acad Sci U S A 112, 15672–15677.

7. Chen, D., Liu, W., Zimmerman, J., Pastor, W.A., Kim, R., Hosohama, L., Ho, J., Aslanyan, M., Gell, J.J., Jacobsen, S.E., et al. (2018). The TFAP2C-Regulated OCT4 Naive Enhancer Is Involved in Human Germline Formation. Cell Rep 25, 3591–3602 e3595.

8. Cornacchia, D., Zhang, C., Zimmer, B., Chung, S.Y., Fan, Y., Soliman, M.A., Tchieu, J., Chambers, S.M., Shah, H., Paull, D., et al. (2019). Lipid Deprivation Induces a Stable, Naive-to-Primed Intermediate State of Pluripotency in Human PSCs. Cell Stem Cell 25, 120–136 e110.

9. Dahan, P., Lu, V., Nguyen, R.M.T., Kennedy, S.A.L., and Teitell, M.A. (2019). Metabolism in pluripotency: Both driver and passenger? J Biol Chem 294, 5420–5429.

10. De Robertis, E.M., and Sasai, Y. (1996). A common plan for dorsoventral patterning in Bilateria. Nature 380, 37–40.

11. Dobin, A., Davis, C.A., Schlesinger, F., Drenkow, J., Zaleski, C., Jha, S., Batut, P., Chaisson, M., and Gingeras, T.R. (2013). STAR: ultrafast universal RNA-seq aligner. Bioinformatics 29, 15–21.

12. Eiraku, M., Watanabe, K., Matsuo-Takasaki, M., Kawada, M., Yonemura, S., Matsumura, M., Wataya, T., Nishiyama, A., Muguruma, K., and Sasai, Y. (2008). Self-organized formation of polarized cortical tissues from ESCs and its active manipulation by extrinsic signals. Cell Stem Cell 3, 519–532.

13. Florio, M., and Huttner, W.B. (2014). Neural progenitors, neurogenesis and the evolution of the neocortex. Development 141, 2182–2194.

14. Garcia-Gonzalo, F.R., and Izpisua Belmonte, J.C. (2008). Albumin-associated lipids regulate human embryonic stem cell self-renewal. PLoS One 3, e1384.

15. Gaujoux, R., and Seoighe, C. (2010). A flexible R package for nonnegative matrix factorization. BMC Bioinformatics 11, 367.

16. Gu, W., Gaeta, X., Sahakyan, A., Chan, A.B., Hong, C.S., Kim, R., Braas, D., Plath, K., Lowry, W.E., and Christofk, H.R. (2016). Glycolytic Metabolism Plays a Functional Role in Regulating Human Pluripotent Stem Cell State. Cell Stem Cell 19, 476–490.

17. Guo, G., von Meyenn, F., Rostovskaya, M., Clarke, J., Dietmann, S., Baker, D., Sahakyan, A., Myers, S., Bertone, P., Reik, W., et al. (2017). Epigenetic resetting of human pluripotency. Development 144, 2748–2763.

18. Hodge, R.D., Bakken, T.E., Miller, J.A., Smith, K.A., Barkan, E.R., Graybuck, L.T., Close, J.L., Long, B., Johansen, N., Penn, O., et al. (2019). Conserved cell types with divergent features in human versus mouse cortex. Nature 573, 61–68.

19. Inman, G.J., Nicolas, F.J., Callahan, J.F., Harling, J.D., Gaster, L.M., Reith, A.D., Laping, N.J., and Hill, C.S. (2002). SB-431542 is a potent and specific inhibitor of transforming growth factor-beta superfamily type I activin receptor-like kinase (ALK) receptors ALK4, ALK5, and ALK7. Mol Pharmacol 62, 65–74.

20. Kadoshima, T., Sakaguchi, H., Nakano, T., Soen, M., Ando, S., Eiraku, M., and Sasai, Y. (2013). Self-organization of axial polarity, inside-out layer pattern, and species-specific progenitor dynamics in human ES cell-derived neocortex. Proc Natl Acad Sci U S A 110, 20284–20289.

21. Kalkan, T., Olova, N., Roode, M., Mulas, C., Lee, H.J., Nett, I., Marks, H., Walker, R., Stunnenberg, H.G., Lilley, K.S., et al. (2017). Tracking the embryonic stem cell transition from ground state pluripotency. Development 144, 1221–1234.

22. Kim, M.H., Kim, M.O., Kim, Y.H., Kim, J.S., and Han, H.J. (2009). Linoleic acid induces mouse embryonic stem cell proliferation via Ca2+/PKC, PI3K/Akt, and MAPKs. Cell Physiol Biochem 23, 53–64.

23. LaMonica, B.E., Lui, J.H., Wang, X., and Kriegstein, A.R. (2012). OSVZ progenitors in the human cortex: an updated perspective on neurodevelopmental disease. Curr Opin Neurobiol 22, 747–753.

24. Lancaster, M.A., Renner, M., Martin, C.A., Wenzel, D., Bicknell, L.S., Hurles, M.E., Homfray, T., Penninger, J.M., Jackson, A.P., and Knoblich, J.A. (2013). Cerebral organoids model human brain development and microcephaly. Nature 501, 373–379.

25. Leek, J.T., Johnson, W.E., Parker, H.S., Jaffe, A.E., and Storey, J.D. (2012). The sva package for removing batch effects and other unwanted variation in high-throughput experiments. Bioinformatics 28, 882–883.

26. Liu, X., Nefzger, C.M., Rossello, F.J., Chen, J., Knaupp, A.S., Firas, J., Ford, E., Pflueger, J., Paynter, J.M., Chy, H.S., et al. (2017). Comprehensive characterization of distinct states of human naive pluripotency generated by reprogramming. Nat Methods 14, 1055–1062.

27. Love, M., Anders, S., and Huber, W. (2014). Differential analysis of count data–the DESeq2 package. Genome Biol 15, 10.1186.

28. Mangale, V.S., Hirokawa, K.E., Satyaki, P.R., Gokulchandran, N., Chikbire, S., Subramanian, L., Shetty, A.S., Martynoga, B., Paul, J., Mai, M.V., et al. (2008). Lhx2 selector activity specifies cortical identity and suppresses hippocampal organizer fate. Science 319, 304–309.

29. Mohammed, H., Hernando-Herraez, I., Savino, A., Scialdone, A., Macaulay, I., Mulas, C., Chandra, T., Voet, T., Dean, W., Nichols, J., et al. (2017). Single-Cell Landscape of Transcriptional Heterogeneity and Cell Fate Decisions during Mouse Early Gastrulation. Cell Rep 20, 1215–1228.

30. Monuki, E.S. (2007). The morphogen signaling network in forebrain development and holoprosencephaly. J Neuropathol Exp Neurol 66, 566–575.

31. Morgani, S., Nichols, J., and Hadjantonakis, A.K. (2017). The many faces of Pluripotency: in vitro adaptations of a continuum of in vivo states. BMC Dev Biol 17, 7.

32. Muguruma, K., Nishiyama, A., Kawakami, H., Hashimoto, K., and Sasai, Y. (2015). Self-organization of polarized cerebellar tissue in 3D culture of human pluripotent stem cells. Cell Rep 10, 537–550.

33. Munoz-Sanjuan, I., and Brivanlou, A.H. (2002). Neural induction, the default model and embryonic stem cells. Nat Rev Neurosci 3, 271–280.

34. Nasu, M., Takata, N., Danjo, T., Sakaguchi, H., Kadoshima, T., Futaki, S., Sekiguchi, K., Eiraku, M., and Sasai, Y. (2012). Robust formation and maintenance of continuous stratified cortical neuroepithelium by laminin-containing matrix in mouse ES cell culture. PLoS One 7, e53024.

35. Nelson, S.B., and Valakh, V. (2015). Excitatory/Inhibitory Balance and Circuit Homeostasis in Autism Spectrum Disorders. Neuron 87, 684–698.

36. Nestler, E.J., and Hyman, S.E. (2010). Animal models of neuropsychiatric disorders. Nat Neurosci 13, 1161–1169.

37. Nguyen, Q.H., Lukowski, S.W., Chiu, H.S., Senabouth, A., Bruxner, T.J.C., Christ, A.N., Palpant, N.J., and Powell, J.E. (2018). Single-cell RNA-seq of human induced pluripotent stem cells reveals cellular heterogeneity and cell state transitions between subpopulations. Genome Res 28, 1053–1066.

38. Ortmann, D., and Vallier, L. (2017). Variability of human pluripotent stem cell lines. Curr Opin Genet Dev 46, 179–185.

39. Pasca, A.M., Sloan, S.A., Clarke, L.E., Tian, Y., Makinson, C.D., Huber, N., Kim, C.H., Park, J.Y., O’Rourke, N.A., Nguyen, K.D., et al. (2015). Functional cortical neurons and astrocytes from human pluripotent stem cells in 3D culture. Nat Methods 12, 671–678.

40. Pastor, W.A., Liu, W., Chen, D., Ho, J., Kim, R., Hunt, T.J., Lukianchikov, A., Liu, X., Polo, J.M., Jacobsen, S.E., et al. (2018). TFAP2C regulates transcription in human naive pluripotency by opening enhancers. Nat Cell Biol 20, 553–564.

41. Qian, X., Nguyen, H.N., Song, M.M., Hadiono, C., Ogden, S.C., Hammack, C., Yao, B., Hamersky, G.R., Jacob, F., Zhong, C., et al. (2016). Brain-Region-Specific Organoids Using Mini-bioreactors for Modeling ZIKV Exposure. Cell 165, 1238–1254.

42. Quadrato, G., Nguyen, T., Macosko, E.Z., Sherwood, J.L., Min Yang, S., Berger, D.R., Maria, N., Scholvin, J., Goldman, M., Kinney, J.P., et al. (2017). Cell diversity and network dynamics in photosensitive human brain organoids. Nature 545, 48–53.

43. Rossin, E.J., Lage, K., Raychaudhuri, S., Xavier, R.J., Tatar, D., Benita, Y., International Inflammatory Bowel Disease Genetics, C., Cotsapas, C., and Daly, M.J. (2011). Proteins encoded in genomic regions associated with immune-mediated disease physically interact and suggest underlying biology. PLoS Genet 7, e1001273.

44. Rostovskaya, M., Stirparo, G.G., and Smith, A. (2019). Capacitation of human naive pluripotent stem cells for multi-lineage differentiation. Development 146, dev172916.

45. Roth, G., and Dicke, U. (2005). Evolution of the brain and intelligence. Trends Cogn Sci 9, 250–257.

46. Sahakyan, A., Kim, R., Chronis, C., Sabri, S., Bonora, G., Theunissen, T.W., Kuoy, E., Langerman, J., Clark, A.T., Jaenisch, R., et al. (2017). Human Naive Pluripotent Stem Cells Model X Chromosome Dampening and X Inactivation. Cell Stem Cell 20, 87–101.

47. Sakaguchi, H., Kadoshima, T., Soen, M., Narii, N., Ishida, Y., Ohgushi, M., Takahashi, J., Eiraku, M., and Sasai, Y. (2015). Generation of functional hippocampal neurons from self-organizing human embryonic stem cell-derived dorsomedial telencephalic tissue. Nat Commun 6, 8896.

48. Samarasinghe, R.A., Miranda, O.A., Mitchell, S., Ferando, I., Watanabe, M., Buth, J.E., Kurdian, A., Golshani, P., Plath, K., Lowry, W.E., et al. (2019). Identification of neural oscillations and epileptiform changes in human brain organoids. bioRxiv, 820183.

49. Seifinejad, A., Tabebordbar, M., Baharvand, H., Boyer, L.A., and Salekdeh, G.H. (2010). Progress and promise towards safe induced pluripotent stem cells for therapy. Stem Cell Rev Rep 6, 297–306.

50. Shiraishi, A., Muguruma, K., and Sasai, Y. (2017). Generation of thalamic neurons from mouse embryonic stem cells. Development 144, 1211–1220.

51. Smith, A. (2017). Formative pluripotency: the executive phase in a developmental continuum. Development 144, 365–373.

52. Takashima, Y., Guo, G., Loos, R., Nichols, J., Ficz, G., Krueger, F., Oxley, D., Santos, F., Clarke, J., Mansfield, W., et al. (2014). Resetting transcription factor control circuitry toward ground-state pluripotency in human. Cell 158, 1254–1269.

53. Theunissen, T.W., Powell, B.E., Wang, H., Mitalipova, M., Faddah, D.A., Reddy, J., Fan, Z.P., Maetzel, D., Ganz, K., and Shi, L. (2014). Systematic identification of culture conditions for induction and maintenance of naive human pluripotency. Cell Stem Cell 15, 471–487.

54. Tidball, A.M., Dang, L.T., Glenn, T.W., Kilbane, E.G., Klarr, D.J., Margolis, J.L., Uhler, M.D., and Parent, J.M. (2017). Rapid Generation of Human Genetic Loss-of-Function iPSC Lines by Simultaneous Reprogramming and Gene Editing. Stem Cell Reports 9, 725–731.

55. Tripathi, S., Pohl, M.O., Zhou, Y., Rodriguez-Frandsen, A., Wang, G., Stein, D.A., Moulton, H.M., DeJesus, P., Che, J., Mulder, L.C., et al. (2015). Meta- and Orthogonal Integration of Influenza “OMICs” Data Defines a Role for UBR4 in Virus Budding. Cell Host Microbe 18, 723–735.

56. van der Worp, H.B., Howells, D.W., Sena, E.S., Porritt, M.J., Rewell, S., O’Collins, V., and Macleod, M.R. (2010). Can animal models of disease reliably inform human studies? PLoS Med 7, e1000245.

57. Velasco, S., Kedaigle, A.J., Simmons, S.K., Nash, A., Rocha, M., Quadrato, G., Paulsen, B., Nguyen, L., Adiconis, X., Regev, A., et al. (2019). Individual brain organoids reproducibly form cell diversity of the human cerebral cortex. Nature 570, 523–527.

58. Wang, L., Zhang, T., Wang, L., Cai, Y., Zhong, X., He, X., Hu, L., Tian, S., Wu, M., Hui, L., et al. (2017). Fatty acid synthesis is critical for stem cell pluripotency via promoting mitochondrial fission. EMBO J 36, 1330–1347.

59. Watanabe, M., Buth, J.E., Vishlaghi, N., de la Torre-Ubieta, L., Taxidis, J., Khakh, B.S., Coppola, G., Pearson, C.A., Yamauchi, K., Gong, D., et al. (2017). Self-Organized Cerebral Organoids with Human-Specific Features Predict Effective Drugs to Combat Zika Virus Infection. Cell Rep 21, 517–532.

60. Watanabe, M., Kang, Y.J., Davies, L.M., Meghpara, S., Lau, K., Chung, C.Y., Kathiriya, J., Hadjantonakis, A.K., and Monuki, E.S. (2012). BMP4 sufficiency to induce choroid plexus epithelial fate from embryonic stem cell-derived neuroepithelial progenitors. J Neurosci 32, 15934–15945.

61. Wataya, T., Ando, S., Muguruma, K., Ikeda, H., Watanabe, K., Eiraku, M., Kawada, M., Takahashi, J., Hashimoto, N., and Sasai, Y. (2008). Minimization of exogenous signals in ES cell culture induces rostral hypothalamic differentiation. Proc Natl Acad Sci U S A 105, 11796–11801.

62. Weinberger, L., Ayyash, M., Novershtern, N., and Hanna, J.H. (2016). Dynamic stem cell states: naive to primed pluripotency in rodents and humans. Nat Rev Mol Cell Biol 17, 155–169.

63. Wickham, H. (2016). ggplot2: elegant graphics for data analysis (Springer).

64. Xu, X., Wells, A.B., O’Brien, D.R., Nehorai, A., and Dougherty, J.D. (2014). Cell type-specific expression analysis to identify putative cellular mechanisms for neurogenetic disorders. J Neurosci 34, 1420–1431.

65. Yoon, S.J., Elahi, L.S., Pasca, A.M., Marton, R.M., Gordon, A., Revah, O., Miura, Y., Walczak, E.M., Holdgate, G.M., Fan, H.C., et al. (2019). Reliability of human cortical organoid generation. Nat Methods 16, 75–78.

66. Youssef, E.A., Berry-Kravis, E., Czech, C., Hagerman, R.J., Hessl, D., Wong, C.Y., Rabbia, M., Deptula, D., John, A., Kinch, R., et al. (2018). Effect of the mGluR5-NAM Basimglurant on Behavior in Adolescents and Adults with Fragile X Syndrome in a Randomized, Double-Blind, Placebo-Controlled Trial: FragXis Phase 2 Results. Neuropsychopharmacology 43, 503–512.

67. Yu, P.B., Deng, D.Y., Lai, C.S., Hong, C.C., Cuny, G.D., Bouxsein, M.L., Hong, D.W., McManus, P.M., Katagiri, T., Sachidanandan, C., et al. (2008). BMP type I receptor inhibition reduces heterotopic [corrected] ossification. Nat Med 14, 1363–1369.

68. Zhang, H., Badur, M.G., Divakaruni, A.S., Parker, S.J., Jager, C., Hiller, K., Murphy, A.N., and Metallo, C.M. (2016). Distinct Metabolic States Can Support Self-Renewal and Lipogenesis in Human Pluripotent Stem Cells under Different Culture Conditions. Cell Rep 16, 1536–1547.

